# Microstructures amplify carotenoid plumage signals in colorful tanagers

**DOI:** 10.1101/799783

**Authors:** Dakota E. McCoy, Allison J. Shultz, Charles Vidoudez, Emma van der Heide, Sunia A. Trauger, David Haig

## Abstract

Red, orange, and yellow carotenoid-colored plumages have been considered honest signals of condition. We comprehensively quantified carotenoid signals in the social, sexually-dimorphic tanager genus *Ramphocelus* using scanning electron microscopy (SEM), finite-difference time-domain (FDTD) optical modeling, liquid chromatography–mass spectrometry (LC-MS), and spectrophotometry. Despite males having significantly more saturated color patches, males and females within a species have equivalent amounts and types of carotenoids. Male, but not female, feathers have elaborate microstructures which amplify color appearance. Expanded barbs enhance color saturation (for the same amount of pigment) by increasing the transmission of optical power through the feather. Dihedral barbules (vertically-angled, strap-shaped barbules) reduce total reflectance to generate “super black” plumage, an optical illusion to enhance nearby color. Dihedral barbules paired with red carotenoid pigment produce “velvet red” plumage. Together, our results suggest that a widely cited index of honesty—carotenoid pigments—cannot fully explain male appearance. We propose that males are selected to evolve amplifiers of honest signals—in this case, microstructures that enhance appearance —that are not necessarily themselves linked to quality.

## Introduction

Why are so many male birds beautiful? To investigate this evolutionary “why,” we study both physical mechanisms of color (pigments and structures) and the evolutionary mechanisms which favor colorful signals over time (selective forces). We must fully understand the physical basis of beautiful traits in order to infer their evolutionary history.

A complete understanding of the physical cause of colorful ornaments may help us determine the relative importance of three overlapping selective pressures within mate choice: species identity, aesthetic beauty, and individual quality. Coloration may facilitate species identification, essential to avoid sterile hybrids and wasted mating efforts (Hill 2015). Beautiful ornaments may reflect arbitrary aesthetic preferences in the choosing sex (Prum 2012), may be maintained through a Fisherian runaway process (Fisher 1999), or may occur as a side effect of selection on another domain such as foraging—termed “sensory bias” (Dawkins and Guilford 1996). Finally, color may be an indicator of individual quality (“honest signaling theory”) due to physiological linkage, resource-trade-offs, or direct/indirect costs (Zahavi 1975; Folstad and Karter 1992; Hill and Montgomerie 1994; Simons et al. 2012) Of the three selective pressures, research to date has largely focused on individual quality, as captured through a large body of literature on honest signaling theory. Frequently, researchers use the physical, pigmentary basis of colorful signals to draw conclusions about honest signaling (e.g., (Velando et al. 2006)).

Carotenoid-pigmented plumages, ranging in hue from yellow to red, have been considered a “textbook example of an honest signal” (Weaver et al. 2018), because carotenoids must be eaten by vertebrates rather than synthesized, may be scarce in nature, and serve important immunological functions (Olson and Owens 1998). However, the relationship between carotenoids, signal appearance, and individual quality is not yet fully understood. Carotenoids are correlated with only some, but not all, individual quality measures (Simons et al. 2012; Koch et al. 2018; Weaver et al. 2018). Carotenoids may be an index of proper metabolic function rather than a costly signal (Weaver et al. 2017). It is often thought that carotenoids are a limiting resource in the immune system; counter to this view, canaries with a mutation knocking out tissue carotenoids show no difference in immune system function (Koch et al. 2018). Nano- and microstructures can alter the appearance of carotenoid-colored organisms (Shawkey and Hill 2005; Iskandar et al. 2016). Is it possible that microstructural elements to carotenoid coloration in birds complicate the link between signal and individual quality, contributing to mixed research results?

The physical basis of color, including microstructural contributions, is not only intrinsically interesting but is also critical to our understanding of sexual selection and evolutionary dynamics. Nanostructures and pigments produce colors through well-understood pathways (Hill and McGraw 2006), but only recently have researchers begun to describe how microstructures can substantially enhance or alter pigmentary colors. Carotenoid pigments produce reds, oranges, and yellows, but microstructures make the colors glossy or matte (Iskandar et al. 2016). The African Emerald Cuckoo *Chrysococcyx cupreus* produces stunning color with milli-scale emerald mirrors (Harvey et al. 2013). Birds-of-paradise (McCoy et al. 2018) and birds from 14 other families (McCoy and Prum 2019) convergently evolved super black plumage with vertically-oriented barbules that enhance melanin-based absorption by multiply scattering light. Outside of birds, many organisms use microstructures to enhance the impact of pigments, from super black peacock spiders (McCoy et al. 2019), stick insects (Maurer et al. 2017), and butterflies (Vukusic et al. 2004; Davis et al. 2020) to flowers with conical epidermal cells that generate richer petal colors (Kay et al. 1981; Gorton and Vogelmann 1996; Gkikas et al. 2015; Wilts et al. 2018). Intriguingly, some researchers have described substantial microstructural variation between male and female birds while overall appearance remains the same (Enbody et al. 2017). To what extent may microstructural, rather than pigmentary, differences explain why so many males are brilliantly colored while many females are drab? The answer may help us determine which selective forces cause so many male birds to be beautiful.

Herein, we aim to better understand the physical basis of color and thus draw inferences about the evolutionary selective pressures which favor colorful signals. We focus on the social, sexually dimorphic *Ramphocelus* tanagers, a useful clade for questions of visual signaling (Burns and Shultz 2012; Shultz and Burns 2017). These tanagers have carotenoid-based coloration ranging from bright yellow to deep velvet red in males, while females are relatively duller (Figure 1). *Ramphocelus* tanager mating behavior has been described for some but not all species; for example, *Ramphocelus costaricensis* has high rates of extra-pair matings (Krueger et al. 2008), and sexual displays have been described in *R. bresilius* (Harris 1987), *R. dimidiatus*, and *R. carbo* (Moynihan 1962).

**Figure 1.**
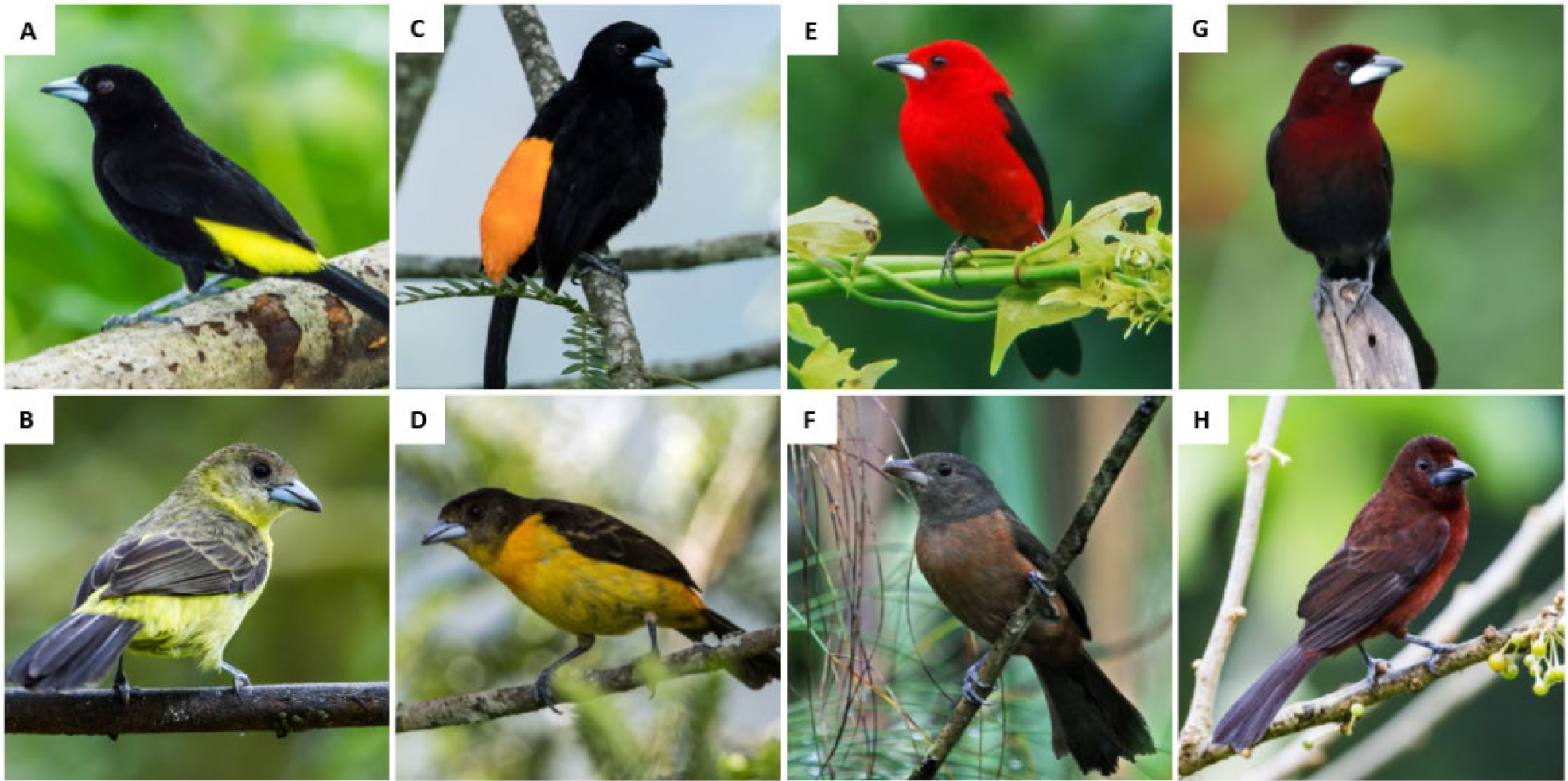
Male *Ramphocelus* tanagers (top row) have more striking carotenoid-based coloration than females (bottom row). A. *R. flammigerus icteronotus* male with vivid yellow and velvety black plumage. B. *R. flammigerus icteronotus* female. C. *R. flammigerus* male with vivid orange and velvety black plumage. D. *R. flammigerus* female. E. *R. bresilius* male with bright red plumage. F. *R. bresilius* female. G. *R. carbo* male with deep, velvety red and black plumage. H. *R. carbo* female. All photos are credit Nick Athanas - www.antpitta.com.

Using scanning electron microscopy (SEM), finite-difference time-domain (FDTD) optical modeling, pigment extraction, liquid chromatography–mass spectrometry (LC-MS), and spectrophotometry, we comprehensively document the physical basis of color in both male and female *Ramphocelus* tanagers. From this, we make inferences about the dynamics of mate choice over evolutionary time. We find evidence that carotenoid-based signals in males are enhanced by substantial microstructural influence. Following substantial past work (Dawkins and Krebs 1978; Krebs and Dawkins 1984; Burk 1988; Dawkins and Guilford 1991; Hill 1994; Parker 2006; McCoy and Haig 2020), we suggest that males, under intense selection to satisfy female testing criteria, evolve “amplifiers” to honest signals (in this case, microstructures that enhance light-pigment interactions).

## Material and Methods

### Specimens

*Ramphocelus* tanager specimens were selected from the Ornithology Collection at the Harvard Museum of Comparative Zoology (MCZ; specimen numbers are listed in Table S1). We selected N=20 total intact male and female specimens, one from each species in the genus *Ramphocelus* plus one subspecies with visually divergent plumage that has had full species status in the past (Griscom 1932): Crimson-collared tanager, *Ramphocelus sanguinolentus;* Masked crimson tanager, *Ramphocelus nigrogularis;* Crimson-backed tanager, *Ramphocelus dimidiatus;* Huallaga tanager, *Ramphocelus melanogaster;* Silver-beaked tanager, *Ramphocelus carbo;* Brazilian tanager, *Ramphocelus bresilius;* Passerini’s tanager, *Ramphocelus passerinii;* Cherrie’s tanager, *Ramphocelus costaricensis;* Flame-rumped tanager, *Ramphocelus flammigerus;* Lemon-rumped tanager, *Ramphocelus flammigerus icteronotus.* Taxonomy is according to the Clements Checklist v.2017 (Clements et al. 2017). We selected only one individual per species because we were primarily focused on interspecific variation, but we confirmed the repeatability of pigment extractions using 3 individuals per species for two species (see below).

We designate *R. flammigerus, R. f. icteronotus, R. passerinii*, and *R. costaricensis* to be the “rumped” tanagers because they form a clade with vivid color restricted to the rump. We designate the clade of tanagers with color on their whole body to be the “whole body” clade (*Ramphocelus nigrogularis, Ramphocelus dimidiatus, Ramphocelus melanogaster, Ramphocelus carbo, Ramphocelus bresilius*). *R. sanguinolentis* is the sister to all others.

### Spectrophotometry

We performed spectrophotometry using an OceanOptics PX2 with a pulsed xenon light source, and OceanOptics USB4000, and an OceanOptics Spectralon White Standard. We used an integration time of 100 ms, averaged 5 scans, and set a boxcar width of 5. We measured three replicates per plumage patch for each of N=20 individuals (male and female from 10 species). For each specimen, we measured the bird at 7 locations: crown, back, rump, dorsal tail, throat, breast, and belly. Using an OceanOptics reflectance probe holder, we measured reflectance for 90° incident light and for 45° incident light. These two angles of incidence allow us to determine the directionality and structural absorption potential of the plumage patches.

We wished to compare saturation of colorful regions and brightness of black regions between males and females. Brightness depends on total reflected light; therefore, we calculated brightness as the integral of the area under the reflectance curve between 300 and 700 nm divided by 400. Saturation depends on the steepness and narrowness of a color peak (Endler 1990); therefore, we calculated an index of saturation by modifying typical full width at half maximum calculations (Eliason et al. 2015). Specifically, we calculated the maximum reflectance, divided that value by two, and calculated the leftwards width of the reflectance curve between the maximum and half-maximum reflectance values. To compare darkness of black regions, we selected the body region of each male and female bird that was darkest (usually backs; see Figure 2). To compare saturation of colorful regions, we selected the body region of each male and female bird that had the highest reflectance (usually rumps; see Figure 2).

**Figure 2.**
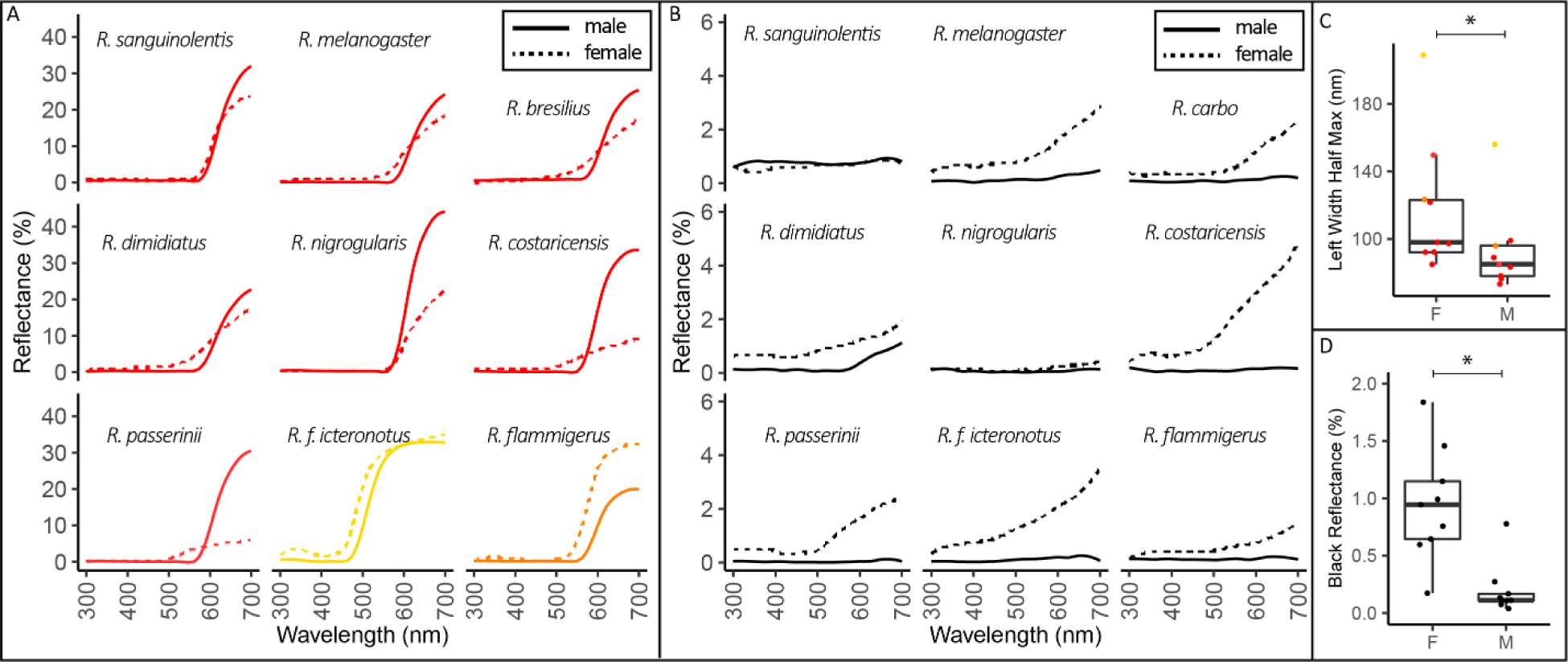
Male Ramphocelus tanagers have significantly more saturated colors and darker blacks than females. Spectrophotometry results for 90° incident light. **A:** Males (solid lines) have brighter and more saturated colors for all species except *R. flammigerus* and *R. f. icteronotus*. We included the body region of each male and female with the max reflectance, which was rump for all species except male *R. sanguinolentis* (crown), male *R. bresilius* (crown), female *R. bresilius* (stomach), female *R. dimidiatus* (stomach), female *R. costaricensis* (breast), and female *R. passerini* (throat). We excluded *R. carbo* from this plot, because it is dark red rather than brilliant; N = 9. **B.** Males (solid lines) have a darker black color, with a flatter reflectance curves, on their backs compared to females (dashed lines) for all species except *R. sanguinolentis*. We measured the black back of each species except *R. dimidiatus*, for which we measured the black throat. We excluded *R. bresilius*, which is all red except for its tail and wings; N = 9. **C.** For a measure of saturation, leftwards width at half maximum reflectance, males had significantly more saturated color than females (phylogenetic two-sample paired t-test; p-value = 0.0072, 95% CI: [13.0, 38.1]). Narrower peaks (i.e., smaller value for left width at half maximum reflectance), indicate a more saturated color. **D.** For a measure of darkness, total integrated reflectance under the curve, males had significantly darker black regions than females (phylogenetic two-sample paired t-test; p-value = 0.0066, 95% CI: [0.38, 1.11]). Complete spectrophotometry results are in Supplemental Data 1. Integration time was 100 ms, and each line represents the average of three replicates within the same plumage patch.

### Mass Spectrometry

We prepared feathers for pigment extraction using a simple mechanical procedure. First, we washed all feathers in hexane and allowed them to air dry. Next, we trimmed off the entirety of colored portions of barbules from three feathers. We carefully weighed them, then placed these barbules with 0.3 mL methanol in a screw cap micro tube in the presence of 10 ceramic beads (2mm). We subjected these tubes to bead beating in a FastPrep FP120 for three cycles, each for 45 seconds at level 6.0. We centrifuged the resulting mixture in an Eppendorf centrifuge 5417R for 5 minutes at 3000 RCF, then transferred the supernatant and dried the barbules under a stream of nitrogen. The samples were resuspended in 100 µL of acetonitrile:methanol (1:1) and transferred to micro inserts in amber glass vials. All samples were kept at - 80 °C until analysis.

Initially, we took 9 feathers per species (three feathers from each of three specimens for two species) to ensure that we had enough material; for all additional species we took 3 feathers from a single patch on a single individual, which proved sufficient. In order to ensure that results were repeatable between individuals of a single species, we extracted pigments from two additional individuals for each of two species—*R. flammigerus* and *R. flammigerus icteronotus*. The pigment profiles were significantly correlated (Figure S3, Linear regression output for *R. flammigerus*: slope = 0.88, SE = 0.066, R^2^ = 0.87, p < 0.0005. Linear regression output for *R. f. icteronotus*: slope = 0.96, SE = 0.054, R^2^ = 0.92, p < 0.0005), indicating that our extraction and characterization procedure was repeatable across individuals of a species. See Statistics section for details. Excluding the repeated measures of *R. flammigerus* and *R. flammigerus icteronotus*, we extracted pigments from N=32 plumage regions in males and females, from 3 feathers per plumage region.

We acquired pigment standards for 8-apo carotenal, Canthaxanthin, Astaxanthin (Sigma Aldrich), Lycopene (Santa Cruz Biotechnology), and Alloxanthin (ChromaDex). Carotenoid contents were analyzed using an Ultimate 3000 LC coupled with a Q-Exactive Plus hybrid quadrupole-orbitrap mass spectrometer (Thermo Fisher Scientific). The LC was fitted with a Kinetex C18 (2.6 µm, 100 Å, 150 mm x 2.1 mm, Phenomenex) column and the mobile phases used were MA: Acetontrile:Methanol 85:15 and MB: 2-propanol. The gradient elution started with 4 min of 5% MB, then to 100% MB in 11 min, followed by 4 min at 100% MB, 1 min to get back to 5% MA and 4 min re-equilibration at 5% MB. The flow was kept constant at 0.185 mL min^-1^, and 10 µL of samples were injected. Electrospray ionization was used in positive ion mode. The mass spectrometer was operated at 70,000 resolving power, scanning the m/z 150-1500 range, with data-dependent (top 5) MS/MS scan at 17500 resolution, 1 m/z isolation and fragmentation at stepped normalized collision energy 15, 35 and 55. The data were first manually analyzed to identify peaks of ions with accurate masses corresponding to potential carotenoids within a 5ppm mass accuracy threshold. The potential carotenoids included both known molecules, and those with a similar molecular formula based on their accurate mass and mass defect. MS/MS fragmentation spectra were then compared with the standards available or to MS/MS database entries within METLIN (-metlin.scripps.edu) to assign identity when standards were available or putative identity based on fragmentation pattern similarity. Lastly, peaks corresponding to carotenoids or carotenoids-candidates were integrated using Quan Browser (Xcalibur, Thermo Fisher Scientific). Pigments were grouped into “families”, based on their accurate mass and relation to standards and fragmentation similarities.

### SEM & Feather Microstructure

In preparation for SEM, we mounted a single feather from each specimen (N=20 feathers) on a pin using a carbon adhesive tab. We sputter-coated the feathers with ∼12 nm of Pt/Pd at a current of 40 mA. We performed scanning electron microscopy on an FESEM Ultra55.

Using ImageJ, we made multiple measurements on each SEM image, including maximum barb width, top-down barb width, barbule width, inter-barbule distance, barb-barbule angle, and barbule length. We also coded feathers according to whether the barbules were strap shaped or cylindrical, and whether the barbules emerged from the plane of the feather (i.e., had “3-D” structure).

### Optical Modelling

To approximate the optical effect of the observed microstructures, we performed finite-difference time-domain (FDTD) simulations using the commercially available software Lumerical FDTD. We constructed idealized two-dimensional feather cross sections for two apparently important microstructural features, as determined by the PCA described below): (i) increasing barbule angle (a purported cause of low-reflectance, super black plumage), (ii) oblong and expanded barb, as is typical in colorful male feathers but not colorful female feathers (a purported enhancer of color). We simulated the interaction of light with these structures to understand the impact of microstructure on plumage appearance. For angled barbules (i), we assessed total reflectance based on structure alone without considering the contribution of melanin. In this manner we can calculate the quantitative decrease in feather reflectance from structure alone. For oblong expanded barbs (ii), we calculated the total optical power transmission through the pigmented portion of the feather. This is a proxy for light-pigment interactions, which strongly influences color: the greater the total optical power transmission through pigment, the greater the pigmentary action. In this manner we can approximate the structural enhancement of pigment due to barb size and shape in males.

FDTD uses the Yee cell method (Taflove and Hagness 2005) to calculate the spatio-temporal electromagnetic field distribution resulting from an initial pulse launched into the simulation domain. For each simulation, a plane wave of light (ranging from 400-700nm) was normally incident (y-direction) on an idealized feather cross-section in the x-y plane. We performed 2D simulations without incorporating pigment because (i) we wished to isolate the effect of structure in a simple, interpretable manner and (ii) the real and imaginary components of carotenoid absorption are not well-understood, as would be required for simulations incorporating pigment. The simulation domain was bounded at the horizontal edges of the y plane by perfectly matched layers (PML) and at the vertical edges by periodic boundary conditions, to simulate a feather plumage. Frequency domain field monitors were placed above and below the structure to collect the reflected and transmitted light, respectively. For our simulations of the oblong, expanded barb, we included a 2D square monitor to capture optical power transmission in a 7.5 x 7.5 µm^2^ square (the purported pigmented region). We used a mesh size of 0.025 x 0.025 µm^2^.

To focus only on the impact of surface reflections and light’s path through the feather after surface diffraction, we extended the bottom feather surface to beyond the bottom vertical PML boundary in the y-plane. This was a measure to eliminate reflections from the underside of the feather for two reasons. First, in life, feathers are arranged in complex stacks in a plumage, which we did not simulate; feathers are flush with other feathers (i.e., the bottom of a feather would be a feather-feather interface, not a feather-air interface). Second, had we included reflections from the underside of a feather, this may over-exaggerate the extent to which microstructure enhances light-pigment interactions.

For our simulations of the male-typical barb vs. female-typical barb, we simulated an average female feather barb based on our measurements: 22.2 µm tall by 18.9 µm wide—and an average male feather barb: 39.8 µm by 27.7 µm. We present the results of one simulation and plot power transmission for 700nm light, but results were similar for other wavelengths of light. For our simulations of angled barbules, we simulated a feather with hemispherical barb (radius 12.5 µm) and barbules 80 µm long varying in angle from 0 to 80°. We present the average and standard deviation reflectance for 15 barbule angles (15 simulations).

### Phylogeny

Several recent studies have examined the phylogeny of *Ramphocelus* tanagers (Hackett 1996; Burns and Racicot 2009; Burns et al. 2014). Unfortunately Burns and Racicot (2009) and Burns et al. (2014) did not include *R. f. icteronotus*, and Hackett (1996) did not include *R. melanogaster* or *R. dimidiatus*, but the two studies combined had *cytochrome b* (cyt b) sequences available for all species. To construct a phylogeny, we downloaded cyt b sequences from NCBI for all *Ramphocelus* species, and six outgroups from the Tachyphoninae clade (*Tachyphonus coronatus, T. rufus, T. phoenicius, Eucometis penicillate, Lanio fulvus, and T. luctuosus*). NCBI accession numbers are available in Table S2. We aligned the sequences with MAFFT v. 7.313 (Katoh and Standley 2013), and constructed the best-scoring maximum likelihood phylogeny with Raxml v. 8.2.10 using the GTRCAT model with 100 bootstrap replicates (Stamatakis 2006, 2014). The resulting phylogeny confirmed the relationships found by previous studies, and so we extracted the monophyletic *Ramphocelus* clade for use in downstream analyses (Figure 3).

**Figure 3:**
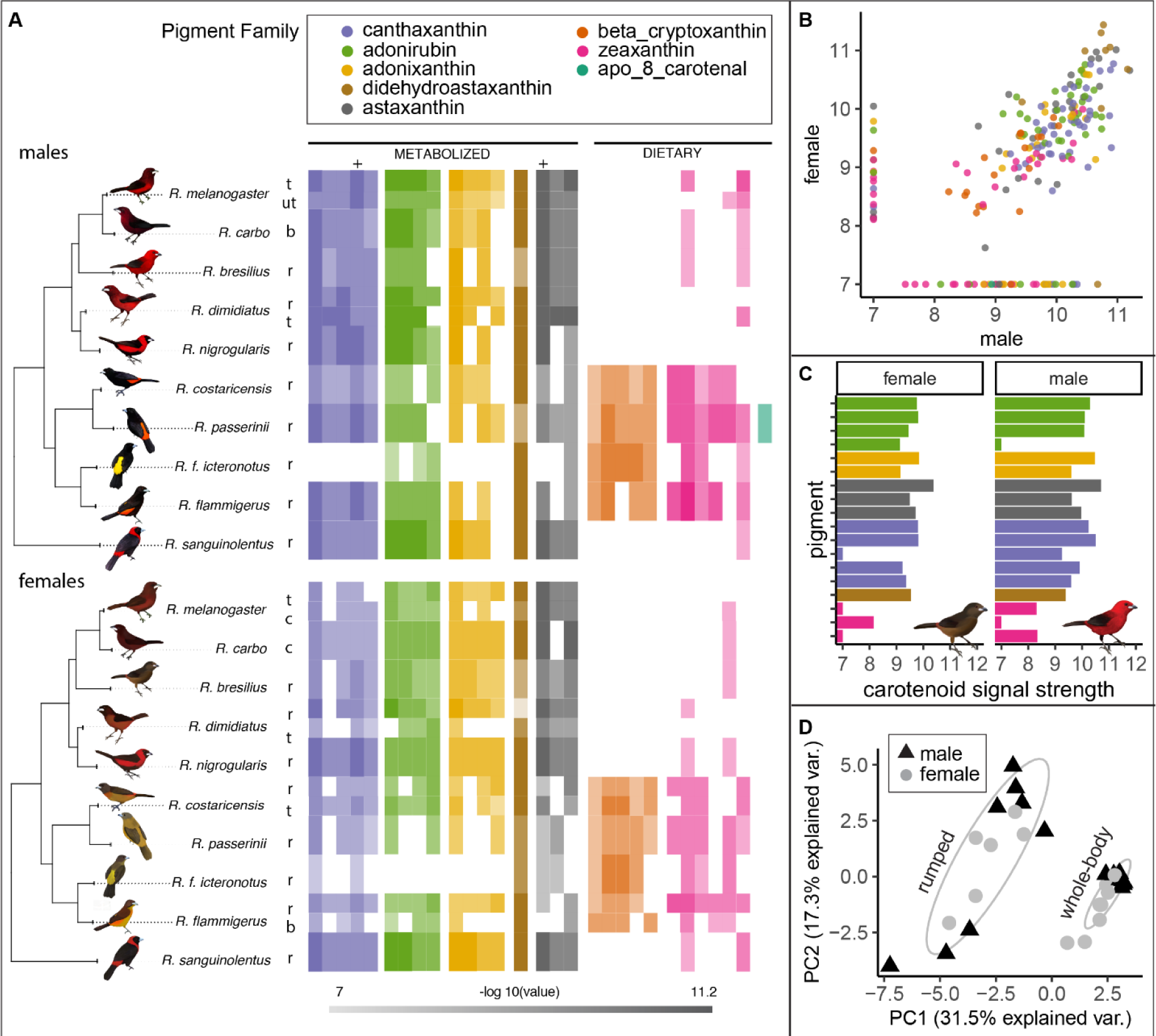
Males and females (within a species) have significantly correlated carotenoid profiles. **A.** Relative log-transformed presence of pigments across birds for males (N_species_ = 10, N_patches_ = 12) and females (N_species_ = 10, N_patches_ = 14). Color saturation indicates normalized signal strength of a given pigment molecule within a bird, where pale is least and rich is most. Relative intensity comparison can therefore only be made within each molecule (column). Some comparisons are possible within each family, because response factors of isomers are expected to be similar. Because standards were not available for all molecules, some molecules were identified based on accurate mass, retention time, MS/MS spectra, and the literature (see Table S4). **B.** Male versus female carotenoid pigments; each point represents the signal strength (a proxy for presence and amount) of a given pigment in both a male and female of one species. All values are log-transformed and normalized. Since 7 was the minimum detectable threshold for our LC-MS setup, we set all values of 0 equal to 7 to better display the data**. C.** Example carotenoid pigment profile for male (top) and female (bottom) *R. bresilius*, demonstrating the similarity. **D.** PCA of log-transformed centered pigments in all patches from all males (N = 10, triangles) and females (N = 10, circles), showing clustering by phylogenetic position (rumped clade separates from whole-body clade). Male and female pigment profiles are significantly correlated (PGLS with pigmentary PC1 scores: coefficient = 0.46, SE = 0.15, p= 0.013). Abbreviations: t: throat; ut: upper throat; r: rump; b: breast; c: chest. Artwork in male silhouettes credit Gabriel Ugueto; female silhouettes are modified by Allison Shultz from original art by Gabriel Ugueto.

### Statistics

To compare saturation and feather brightness in males and females, we performed phylogenetically controlled two-sample t-tests in R, using phyl.pairedttest from package phytools v. 0.6-44 (Revell 2012, 2013).

We performed principal component analyses (PCAs) on microstructure and pigment levels in males and females, using both standard and phylogenetic PCA methods in the software program R, version 3.4.3. We log transformed and centered all data. For standard PCAs we used function prcomp in R, both centering and scaling the data, on measurements of feather microstructure (Table S3) and pigment signal strengths for all feathers from all species, both males and females. PC1 and PC2 were extracted and used for visualizations; PC1 was used to assess correlations between males and females.

For phylogenetic PCAs, we used function phyl.pca in the R package phytools (Revell 2012), with a lambda method of correlation, and PCA mode “cov” (covariance). For phylogenetic PCAs, we had to reduce the character matrix, so we summed the pigment signals within each pigment family to generate 8, rather than 29, individual characters. Phylogenetic PCAs cannot incorporate more than one measurement per species, so we had to run separate phylogenetic PCAs for males and females.

We tested for correlations between the proportions of carotenoids detected (carotenoid signal, as peak area) in two replicates each of two species, *R. flammigerus* and *R. f. icteronotus*. First, for each individual, we normalized the signal for each carotenoid to account for variation in overall amounts of carotenoid detected (versus proportion) by dividing the signal of each carotenoid by the sample weight and then by the maximum signal detected for that individual. We then tested for a correlation between the two individuals of each species separately with a linear model. We performed the same procedure to test for correlations between the upper and lower throat of male *R. melanogaster* and the throat and rump of male *R. dimidiatus*.

We also tested for correlations among the proportions of male and female carotenoids and male and female feather structure. Because it is not possible to test more than one character per species at a time in a phylogenetic framework, we decided to test for correlations between male and female measurements using the PCA scores, which captured a large proportion of the variance. For both carotenoids and structure, we tested PC1 and PC2, and used phylogenetic generalized least squares (PGLS; (Grafen 1989; Martins and Hansen 1997)) with a Brownian motion model to account for phylogeny. We could not use PGLS for multiple patches when measured, so we randomly selected one patch for each species. Analyses using the alternative patch were qualitatively similar, but not presented here. Results are also robust to PGLS model choice for phylogeny transformation. Note that these PC scores were taken from the non-phylogenetic PCA, and thus are directly comparable between males and females (both of which were included in the PCA).

Finally, we reconstructed the evolutionary history of carotenoid evolution for each pigment family for males and females. Because there might have been slight variations in isoform for different species, we decided to focus on carotenoid families as the most biologically meaningful variables. First, for each pigment family, we summed all isoforms for males and females, and log-transformed them (base 10). We removed any species that were missing data for both males and female for that family. We did the reconstruction with the contMap function in phytools v. 0.6-44 (Revell 2012, 2013), which uses maximum likelihood to estimate states at internal nodes and interpolate these states along internal branches (Felsenstein 1985).

## Results

### Male Ramphocelus tanagers have significantly more saturated colors and darker blacks than females

Spectrophotometry revealed a wide range of yellows, oranges, and reds in males and females, adjacent to blacks (Figure 2, complete results in Supplementary Data 1). Vivid, highly saturated color patches in males typically reflected almost no light for short wavelengths before sloping sharply upwards to reflect long wavelengths in the yellow-orange-red space (Figure 2A). In contrast, females of most species had colorful patches with a relatively more gradual upward slope, a greater-than-0 reflectance over a broad range of wavelengths, and a relatively lower peak reflectance (Figure 2A). Males from multiple species had extremely low-reflectance black plumage, with broadband flat reflectance below 0.5%, in areas adjacent to bright color patches (Figure 2B). Females had typical black plumage reflectances, with a slight increase in reflectance at higher wavelengths (Figure 2B).

Males had significantly more saturated color than females (phylogenetic two-sample paired t-test; p-value = 0.0072, 95% CI: [13.0, 38.1]) based on a measure of saturation, leftwards width at half maximum reflectance (Figure 2C). That is, they had narrower peaks. Males had significantly darker black regions than females (phylogenetic two-sample paired t-test; p-value = 0.0066, 95% CI: [0.38, 1.11]), where darkness was measured as total integrated reflectance under the curve (Figure 2D). One species, *R. carbo*, has dark red velvet color on its whole body. Spectrophotometry showed that this plumage reflected very little light (Supplementary Data 1) in a manner similar to the velvet black feathers in other species (Figure 2B).

In general, many of the colored feathers were strong directional reflectors, such that reflected light differed in quantity (percent total reflectance) from 90° incident light to 45° incident light (Supplementary Data 1). For example, the red velvet patches on *R. carbo* increased their reflectance from ∼0-1% to roughly 5-7% when the angle of light incidence changed from 90° to 45°. This characteristic, reflecting more light at a lower angle of incidence, is a hallmark of feathers with microstructures that impact absorbance and reflection (McCoy et al. 2018).

### Males and females (within a species) have significantly correlated carotenoid profiles

We identified 29 distinct molecules that matched the absorption spectra of carotenoids. These fell into 8 groups by monoisotopic molecular mass, with 1-6 different molecules in each pigment group, which we refer to as pigment families (Table S4). We mapped the relative abundances of all described molecules onto a tree for both males and females (Figure 3A), performed ancestral state reconstructions for each pigment family (Figure S1), and mapped all identified pigments onto a metabolic network that shows how different molecules can be modified within the body (Figure S2).

Five of the molecules matched our purchased pigment standards and could be conclusively identified. For the remaining pigments, the molecule’s identity was inferred based on accurate mass, retention time, MS/MS spectra, and pigments commonly found in bird feathers as described in the literature. For example, it is difficult to distinguish between the two dietary carotenoids lutein and zeaxanthin. Using the metabolic networks presented in Morrison and Badyaev (2016) and LaFountain et al. (2015), we identify the molecules with the accurate mass 568.43 as zeaxanthin because none of the common avian derivates from lutein were found in any of our samples, while many derivates of zeaxanthin (and related metabolites) were found. If some or all of the detected molecules with this mass were lutein, it would not change the conclusions herein, because both are dietary pigments.

We found that dietary pigments zeaxanthin, β-cryptoxanthin, and apo-8-carotenal were found primarily in the rumped tanager clade (Figure 3A, blue stars in Figure S2) while the whole-body tanagers and *R. sanguinolentis* had primarily metabolized carotenoids. The “rumped” tanager clade had less metabolized pigments than the “whole-body” clade and *R. sanguinolentis* (Figure 3A). This was affirmed through ancestral state reconstructions (Figure S1). These mappings also showed qualitative concordance between male and female pigment profiles within a species, and qualitative concordance between pigment profiles of different regions within a bird (Figure 3).

We found that pigment profiles were significantly correlated between males and females within species (Figure 4A; PGLS with pigmentary PC1 scores: coefficient = 0.46, SE = 0.15, p= 0.013). In other words, we performed a PCA of all pigment signal strengths in both males and females to generate a single PC1 score that represented pigment content (Figure 6A). We then assessed the correlation between male pigmentary PC1 scores and female pigmentary PC1 scores, controlling for phylogeny with PGLS, to determine whether males and females have similar carotenoid pigment profiles. To visualize this result, we plotted the relative signal strengths (a proxy for amount) of each pigment in feathers from a male and female of the same species (Figure 3B), along with example pigment profiles from male and female *R. bresilius* (Figure 3C). These plots of the raw data (Figure 3A-C) qualitatively support our quantitative conclusions based on PC1: male and female tanagers within a species have concordant pigment profiles.

**Figure 4:**
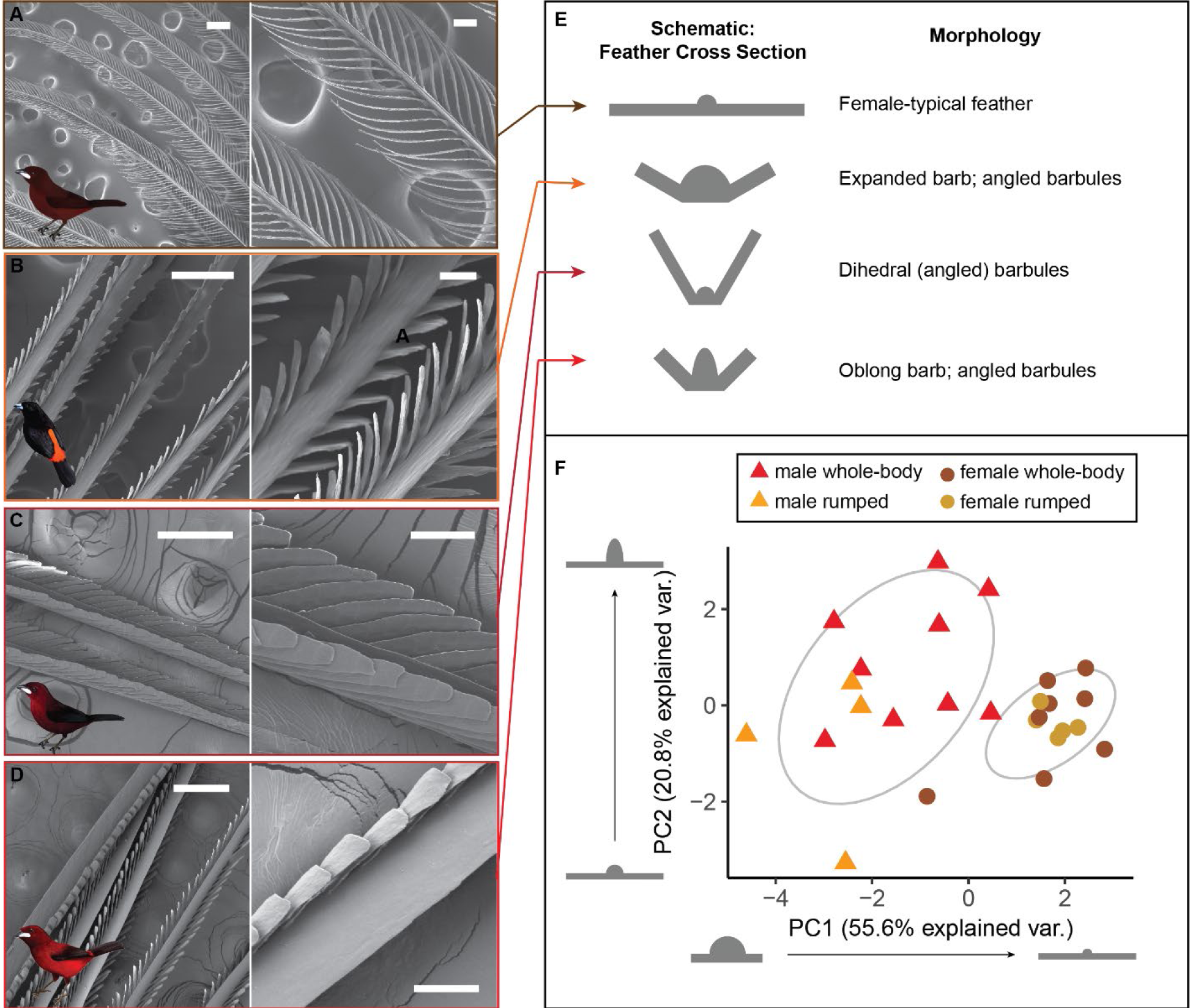
Males, but not females, have diverse and elaborate feather microstructure. **A.** Female *R. carbo* red chest feather with typical simple morphology. **B:** Male *R. passerinii* bright orange rump feather with expanded barb and strap-shaped barbules. **C.** Male *R. carbo* velvet red-black back feather with dihedral barbules. **D:** Male *R. dimidiatus* shiny red rump feather with expanded, oblong barb and strap-shaped barbules. Scale bars in left column are 200 µm; scale bars in right column (B, D, F, H) are 50 µm. Artwork in inset silhouettes credit Gabriel Ugueto. **E.** Schematic illustrations of idealized cross-sections of each feather type in A-D. **F.** PCA of log-transformed centered microstructural measurements for all patches from all males and all females, showing that males separate from females. Male and female microstructures are not correlated (PGLS with microstructure PC1 scores: coefficient = -0.023, SE = 0.17, p= 0.90). This PCA does not take into account differences in three-dimensional structure, including dihedral barbule morphology; see Table 1.

**Figure 5:**
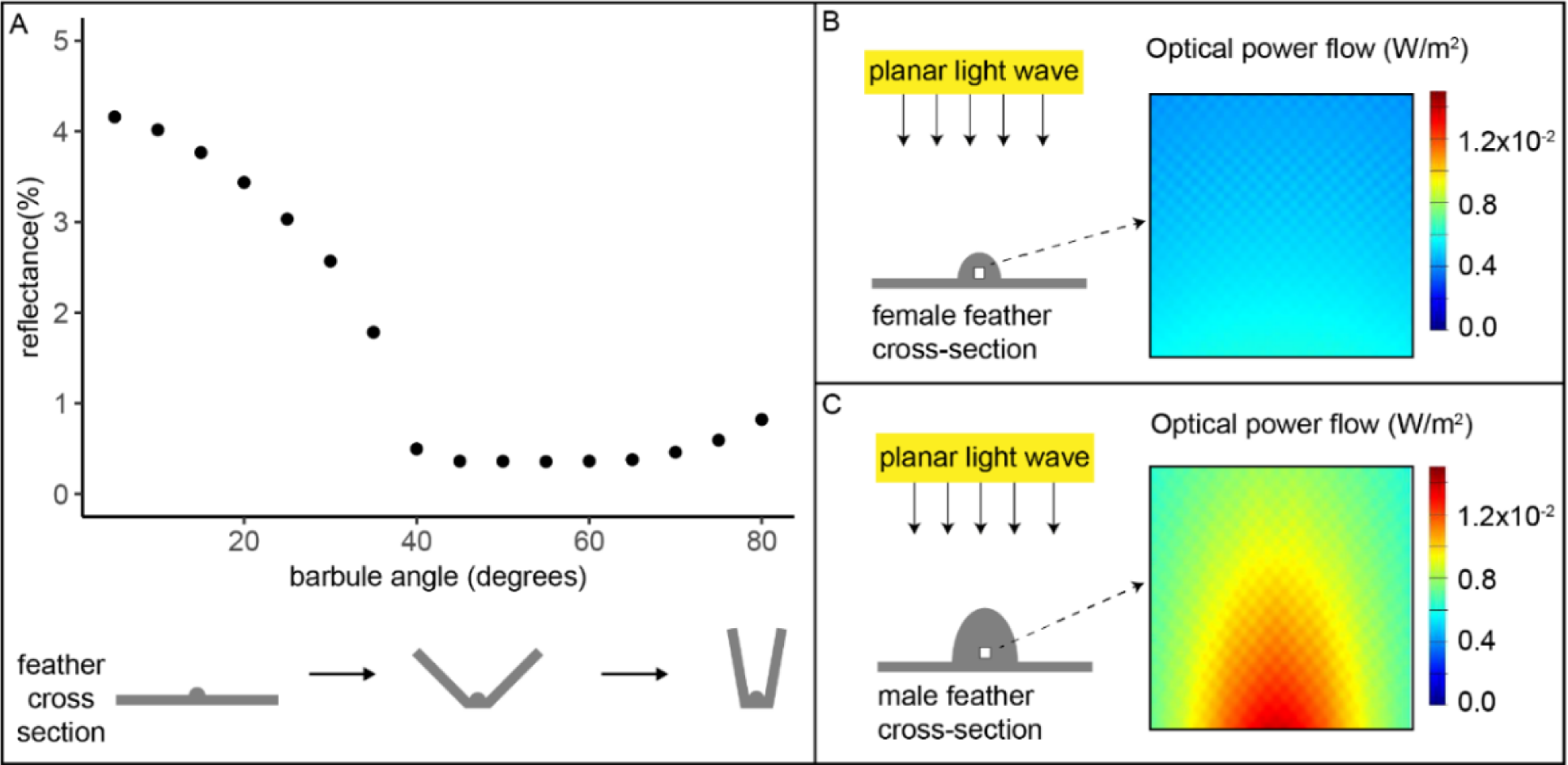
Optical simulations show that male-typical microstructures can (i) enhance blackness and (ii) increase color saturation. All simulations are finite-difference time-domain (FDTD) simulations on 2D idealized feather cross-sections, simulating structural effects only without accounting for pigmentary absorption. **A:** A male-typical black feather, with angled barbules, has structurally enhanced blackness compared to a female-typical feather with flat barbules. As the barbule angle increases out of the plane of the feather, percent reflectance arising from the keratin structure alone decreases from 4% to 0.5%. We did not simulate absorption effects of melanin, but instead we isolated the effect of structure alone. **B:** For a female-typical colored feather, optical power flow through the purported pigmented area (white square) ranges from 0.004-0.005 W/m^2^. **C**: For a male-typical colored feather, an oblong, expanded central barb increases optical power transmission through the purported pigmented area (white square; 0.005-0.016 W/m^2^). This enhanced power transmission through the pigmented region enhances pigmentary activity for the same amount of incident light. For A, each dot is the average of 3 simulations (and a scale bar is plotted although not visible because all standard deviations were < 0.035). For B-C, optical power transmission is shown for 700nm light and for a single simulation, but the results are similar for other wavelengths of light and for repeated simulations.

**Figure 6:**
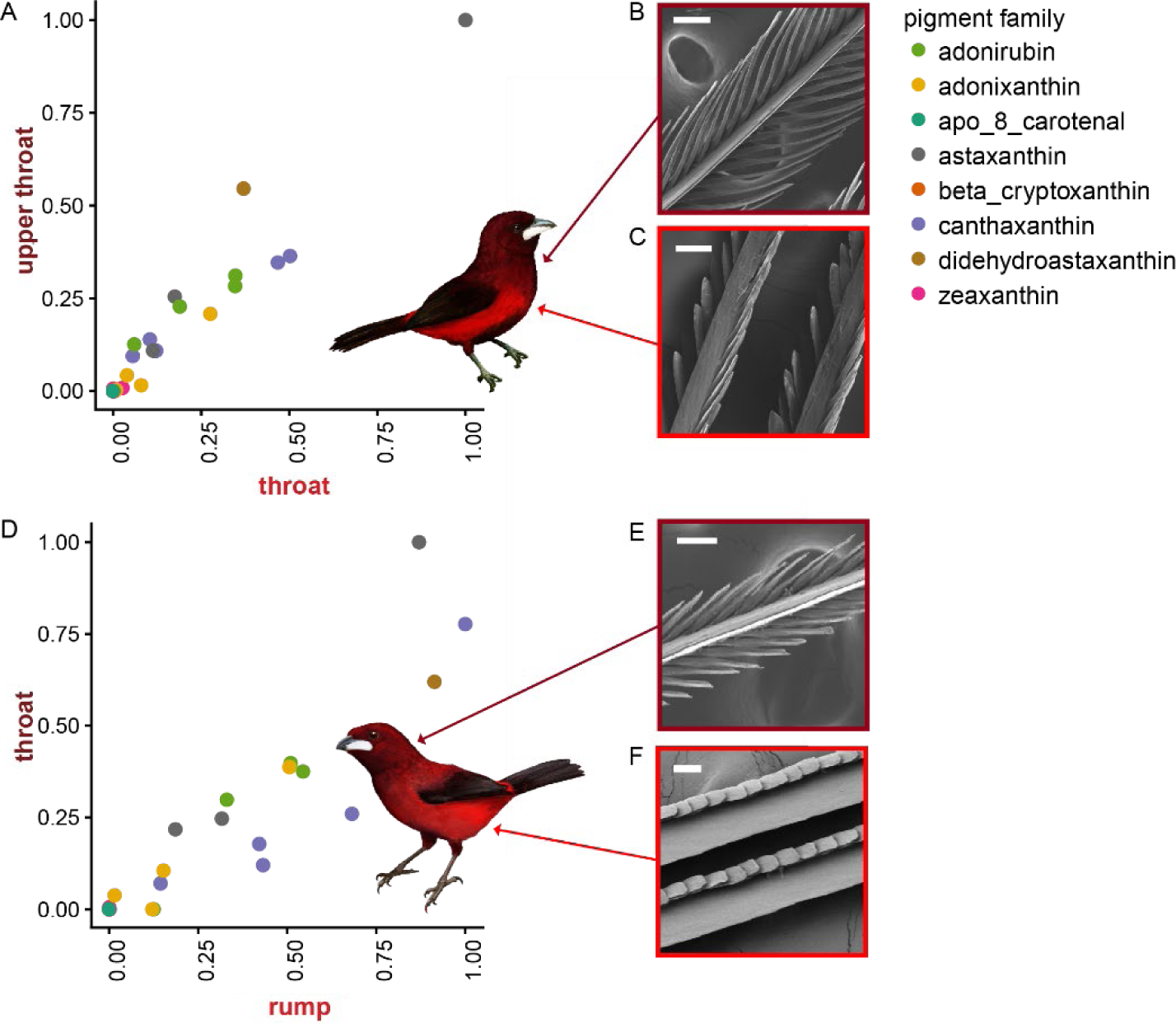
Microstructure, not carotenoids, differs between bright and dark red patches within a bird. **A:** *R. melanogaster* dark velvet red throat versus brighter red lower throat. Pigments are highly correlated between regions: slope = 0.99, SE = 0.051, R^2^ = 0.93, p < 0.0005. **B-C:** *R. melanogaster* with SEM images of dark red throat feather (top) compared to brighter red lower throat feather (bottom). **D:** Pigment profiles of *R. dimidiatus* dark velvet throat versus bright red rump, where each point represents the signal strength for one pigment molecule. Pigments are highly correlated between regions: slope = 0.76, SE = 0.061, R^2^ = 0.85, p < 0.0005. **E-F:** *R. dimidiatus* with SEM images of dark red throat feather (top) compared to bright red rump feather (bottom). All values are normalized such that the largest value (strongest signal) was set equal to one. Scale bars are 50 µm. Artwork in bird silhouettes credit Gabriel Ugueto.

A PCA of carotenoid pigments in all feathers from both males and females demonstrated that birds do not separate by sex, but instead separate by phylogeny (rumped clade separate from whole-body clade; Figure 3D; PC1 31.5%, PC2 17.3%). These results are concordant with separate phylogenetic PCAs of males and females (Figure S4, Table S6).

**Table 1:**
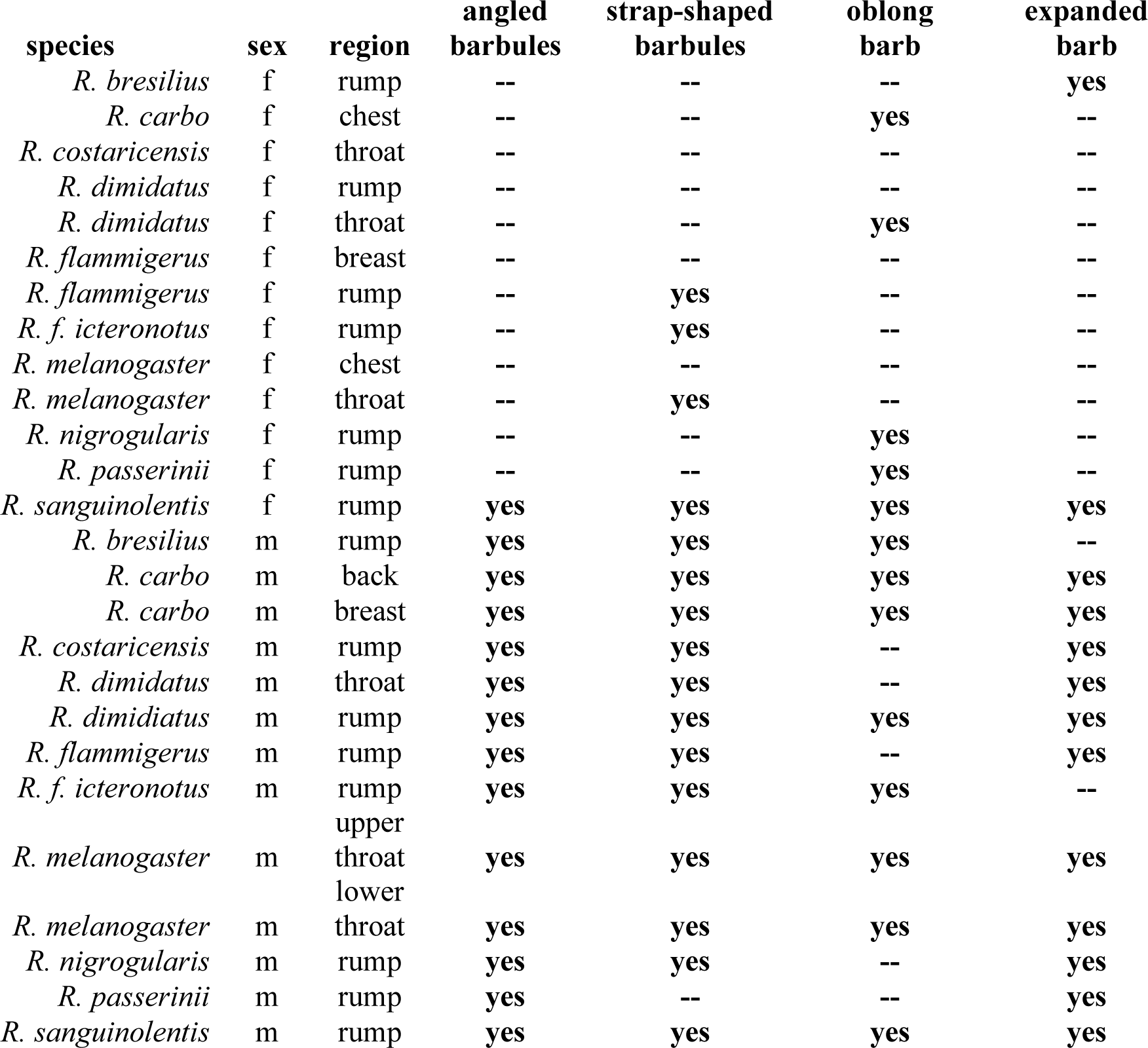
Males, but not females, have diverse and elaborate feather microstructure. Male *Ramphocelus* tanagers have many atypical microstructural features compared to females. “3D barbules” refers to whether or not barbules extended upwards from the plane of the feather; “strap-shaped barbules” refers to whether barbules were flattened rather than classically cylindrical; “oblong barb” refers to whether barbs were taller than they were wide (numerical threshold: > median (max barb width)/(top-down barb width)); wide barb refers to whether the barb was expanded (numerical threshold: > median barb width value). Complete numerical measurements can be found in Table S3.

### Males, but not females, have diverse and elaborate feather microstructure

Female *Ramphocelus* tanager feathers were mostly of standard microstructural appearance (McCoy and Prum 2019). That is, feather microstructure usually looks the same as feather macrostructure, with simple cylindrical barbules extending from the central cylindrical barb, all in a single plane (Figure 4A, Table 1, Table S3). The only female to exhibit broad variation was *R. sanguinolentis* (Table 1, Figure S5, Table S3).

In contrast, male *Ramphocelus* tanagers exhibited wide variation in feather microstructure (Figure 4, Table 1, Table S3, Figure S5), including widely expanded barbs, oblong barbs, strap-shaped barbules (rather than cylindrical), and angled barbules that projected from the plane of the feather (Table 1). Among the angled barbules, we observed a dihedral morphology in velvety black and velvety red feathers as described previously for super black feathers of *R. flammigerus* (McCoy and Prum 2019). Two of these unusual male morphological traits, which we assess through optical simulations, deserve special mention (Figure 4):

#### Dihedral

In the dark red crown of *R. carbo*, feathers have densely packed, strap-shaped barbules pointing upwards out of the plane of the feather to form a dihedral structure, while the central barb was shaped like a razor or narrow triangular prism (Figure 4C). *Ramphocelus dimidiatus* shared the “red velvet” morphology present in *R. carbo* in its upper throat feathers, which are a dark velvety red color (Figure S5). Both are reminiscent of the structurally absorbing “super black” feather morphology present in *Ramphocelus flammigerus* (McCoy and Prum 2019). Similar morphology, to a lesser degree, was observed in *R. melanogaster* dark velvet throat feathers and in red feathers of both male and female *R. sanguinolentis* (Figure S5G,J). To the human eye, dihedral feather structure generates a velvety appearance.

#### Expanded barb

Multiple species had expanded barbs, including the broad and roughly cylindrical barbs of the rump feathers of *R. flammigerus* (e.g., Figure 4B,D, *R. nigrogularis*, and *R. passerinii*). These feathers also had flatter, strap-shaped barbules shorter in length than that of females but wider in width. Additionally, the rump and body feathers of *R. dimidiatus* are vivid, glittery red feathers with a more extremely expanded central barb with a featureless surface (Figure 4D). They also have upward-slanting, densely packed, strap-shaped barbules (Figure 4D). Together, this generates (to the human eye) strong specular reflection at an angle, so that the bird glitters when rotated in the hand.

A PCA of feather microstructure measurements demonstrated that males and female cluster separately and are not correlated (Figure 4F; PC1 55.6%, PC2 20.8%; PGLS with microstructure PC1 scores: coefficient = -0.023, SE = 0.17, p= 0.90); a phylogenetic PCA was not possible to compare males and females, because such analyses can consider only one value per species, but we performed sex-specific phylogenetic PCAs of one patch per species and observed some clustering by clade for males (whole-body versus rumped, Figure S4) and that females cluster tightly with the exception of outliers *R. sanguinolentis* and *R. bresilius*, both of which are females with wider-than-median barbs (Table 1). *R. sanguinolentis* is also the female with the most male-like scores in the regular PCA (Figure 4F). Barb width, barbule width, barb oblongness, and interbarbule distance were primary axes along which males and females separated (Table S5). No species-specific or region-specific (e.g., rump feather versus chest feather) clustering was observed in these microstructural measurements (Figure S4).

### Optical simulations show that male-typical microstructures can enhance blackness and colorfulness

Using 2D finite-difference time-domain (FDTD) simulations of idealized feather cross-sections, we isolated the effect of two male-typical structure on reflectance: dihedral barbules and expanded barb. We selected these two features because they were associated with (i) within-bird color changes from bright, saturated red to velvety red (Figure 6) and (ii) male-female differences in color appearance within a species.

We found that “dihedral” barbules (McCoy and Prum 2019), projecting out of the plane of the feather and associated with super blacks and velvet reds in our study species, contribute to a lower reflectance (Figure 5A). Total reflectance based on light-keratin interactions alone, without considering the contribution of melanin or carotenoids, drops from ∼4% to 0.5% as barbules increase in angle from 0° to 80°. That is, the dihedral feather morphology in super black regions of these tanagers are made blacker in part due to structure. Likewise, the velvet red feathers in *R. carbo* and *R. dimidiatus* are made darker and velvety by structure.

Further, we found that male-typical expanded feather barbs, which are wider than female barbs in 80% of species and more oblong than female barbs in 60% of our species (Table 1), substantially enhance optical power flow through the pigmented portion of the feather (Figure 5B-C). In other words, this male-typical barbule shape causes greater light-pigment interactions, even though females and males have the same types of pigment (Figure 3). Therefore, the structure may contribute to a richer, more saturated color in the same manner as flower petals’ conical epidermal cells (Gkikas et al. 2015).

### Microstructure, not carotenoids, differs between bright and dark red patches within a bird

To better assess the relative contribution of microstructure and pigment, we compared within-bird pigment profiles of shiny saturated red patches versus dark velvety red patches for both *R. melanogaster* and *R. dimidiatus* (Figure 6). For each bird, these two patches had significantly correlated pigment profiles at equal levels for *R. melanogaster* and with slightly more pigment in the bright red region for *R. dimidiatus* (Figure 6A,D; Regression output for *R. melanogaster*: slope = 0.99, SE = 0.051, R^2^ = 0.93, p < 0.0005. **B:** Regression output for *R. dimidiatus*: slope = 0.76, SE = 0.061, R^2^ = 0.85, p < 0.0005.) However, the SEM photos revealed large differences in feather microstructure between velvety and bright patches (Figure 6B,C,E,F). Expanded barbs were associated with bright saturated color while vertically-angled or dihedral barbules were associated with velvety color. Within-bird color varies according to microstructure, not carotenoids, which supports the results of our optical modeling.

## Discussion

### Summary

Why are so many male birds beautiful? Our study directly addressed a proximate, physical “why” by quantifying the determinants of color. Colorful male and significantly more drab female *Ramphocelus* (Figures 1-2) have the same amounts and types of carotenoid pigments (Figure 3). However, unusual microstructures in male (but not female) feathers augment male appearance in two ways (Figures 4-5). First, wide, oblong barbs in bright red and orange male feathers enhance light-pigment interactions to create a more vivid, saturated color from structure alone without requiring more pigment. Second, dihedral barbules produce (i) low-reflectance super black near colorful patches, an optical illusion for color emphasis (McCoy et al. 2018, 2019; McCoy and Prum 2019)), and (ii) velvety red feathers adjacent to brilliantly reflective beaks (Figure 1). We observe that structures contribute significantly to color signal appearance in males, rather than pigment alone.

But *why* are so many male birds beautiful? A deeper evolutionary “why” asks for the history of selective forces that produce beauty. Carotenoid-based coloration has often been invoked as a “textbook” (Weaver et al. 2018) or “classic” (Grether et al. 2001) example of honest signaling. Carotenoids are thought to be honest because they are rare, are costly, require a trade-off, and/or are an index of proper metabolic function (Hill and Montgomerie 1994; Olson and Owens 1998; Faivre et al. 2003; Saks et al. 2003; Simons et al. 2012; Weaver et al. 2017, 2018; Koch et al. 2018). We cast doubt on the carotenoids-as-*costly*-signals theory by showing that males and females have equivalent carotenoid pigment profiles. More broadly, can carotenoid-colored signals in *Ramphocelus* be considered “honest,” given that microstructures significantly alter appearance? Why do males have such diverse and influential microstructures at all? We propose that carotenoid coloration may have originated as an honest signal of quality, but males have been selected to amplify their appearance by microstructural enhancers that are not themselves necessarily linked to quality. We do not suggest that signals have no indicator value, but rather that diverse microstructures in males suggest an evolutionary arms race between female preference and male appearance. We discuss this “proxy treadmill” interpretation (McCoy and Haig 2020), and other possibilities, below.

### Pigments: males and females within a species have the same carotenoid profiles

We found a surprising 29 distinct carotenoid molecules in the plumages of *Ramphocelus* tanagers (Figure 3), some of which are dietary carotenoids-consumed and directly deposited into feathers-while others are metabolized. If carotenoids were present in feathers as an honest (because costly) signal, it would be surprising to see the same levels in females (the choosing sex) as in males (the displaying sex). Unexpectedly, males and females of the same species had similar carotenoid profiles in feathers, with similar levels and types in both sexes (Figure 3). A correlation between male and female signals despite dimorphism can result from general constraint and opportunities of a shared genome (Lassance 2009). For example, female club-winged manakins (*Machaeropterus deliciosus*, Pipridae) select males who have altered wing feathers and bones in order to produce musical sound; oddly, the females have modified bones as well, which may hurt their adaptive survival (Prum 2017). Male and female tanagers have been able to evolve different feather structures. Therefore, it is difficult to argue that the presence of carotenoids in feathers is costly, is favored in males because it is costly, but females who gain no benefit therefrom have been unable to exclude carotenoids from their feathers. However, it is still possible that carotenoids are an index of metabolic function, rather than an example of a costly investment or trade-off (Weaver et al. 2017). It is also possible that sexual selection is acting on both males and females. Further work comparing carotenoid presence across many male museum specimens, as well as color signal quality, may help clarify this question.

It is worth dedicating a few sentences to the significance of dietary versus metabolized carotenoids. Importantly, carotenoids are consumed in a yellow (dietary) form and then metabolized within vertebrate bodies to become immunologically useful molecules that are redder in color (Weaver et al. 2018)w. A recent meta-analysis showed that metabolized, rather than dietary, carotenoids drive correlations between male appearance and health (Weaver et al. 2018). Perhaps red metabolized carotenoids are thus a signal of a functioning metabolism. However, in *Ramphocelus* it appears that *R. f. icternonotus* and *R. flammigerus* evolved to use dietary carotenoids (yellow & orange) from a likely ancestral state of metabolized (red) carotenoids (Figure 3, Figure S1; this is the most parsimonious explanation, particularly given that the *Ramphocelus* clade has red-colored sister species (Burns and Racicot 2009)). In fringillid finches, a model system for plumage evolution, there are no documented reversions from metabolized to dietary pigments (Ligon et al. 2016). New world blackbirds (Icteridae) convergently evolved red plumage from yellow six times but never in the reverse direction (Friedman et al. 2011). Reversions to yellow seem rare (i.e., less likely than gains of red), and our tree may indicate two independent origins of red rather than one loss. However, our data demonstrates that yellow and orange birds do have some red metabolized pigments (Figure 3), just in lower quantities than in the red birds, and are thus capable of the metabolic processes that convert dietary to metabolized pigments. The possible reversion from red to yellow and orange documented herein could be evidence that *Ramphocelus* males were not selected to display metabolically indicative or costly traits through carotenoid pigmentation. Further detailed chemical analyses of the pigments which are differentially expressed between males and females (Figure 3) would be valuable.

Further research should be done on the behavioural ecology of *Ramphocelus* to clarify female preferences, particularly regarding dietary (yellow/orange) versus metabolized (orange/red) pigment signals. In a hybrid zone, Morales-Rozo and colleagues demonstrate that the pigment-poor, yellow *R. f. icteronotus* outcompetes the comparatively pigment-rich, orange *R. flammigerus* in areas where both male phenotypes occur (i.e., where females have a choice)(Morales-Rozo et al. 2017). Specific mate-choice experiments would be needed to demonstrate this conclusively, but if females do indeed prefer this less-elaborate, less-costly trait, this aligns well with Hill’s (1994) criteria for an arbitrary, rather than honest, model of trait evolution: Hill predicted that under honest advertisement models, “in lineages in which a less elaborate form of a trait is derived from a more elaborate form, females should show a preference for the primitive, extreme expression of the trait” (Hill 1994).

In addition to the pigments we document, the relative amounts of other pigment types, such as melanin, or internal nanostructural changes likely also play a role in the different appearances of males and females. This is a worthwhile focus of future studies, as well as the physiological cost or importance of other pigments and nanostructures.

### Structures: microstructures amplify plumage appearance in males

Feathers in *Ramphocelus* males, but not females, vary remarkably in microscopic structure (Table 1, Figure 4, Figure S5). Relative to the typically-proportioned female feathers, many males have extremely expanded barbs and strap-shaped, upward-angled barbules. It is microstructure, not carotenoid pigment, that differs between males and females (Figure 3-4). With optical modelling, we show that expanded barbs enhance color and upward-angled barbules generate velvety red and super black (Figure 5)—a deep black which frames and amplifies nearby color. That is, structures alone enhance appearance independently of pigment content.

Oblong, expanded central barbs enhance color because they increase the optical power transmitted through the feather (Figure 5B-C). Male feathers have more oblong and more expanded barbs (Table 1). More light interacts with the same amount of pigment to generate a more saturated, vivid color in males (Figures 1-2). This is analogous to a pigmented solution in a glass test tube, which appears more-richly-colored from certain angles due to the differences in path length of light through the solution.

Upward-angled barbs in a dihedral arrangement generate a velvety, super black appearance (Figure 5A)—an optical illusion that enhances nearby colors. These “super black” plumage patches reflect almost no 90° incident light and have a broadband, featureless reflectance spectrum in contrast to a rise in reflectance at higher wavelengths characteristic of melanin pigmentary reflection alone (Figure 2B). Normally, vertebrates use white gleams (mirror-like specular reflections) to control for the quantity of ambient light, thus accurately perceiving color across many lighting conditions. Super black enhances the perceived brilliance of adjacent colors by removing nearly all specular reflection, thus influencing the innate control capacities of vertebrate eyes and brains (Kreezer 1930; Brainard et al. 1993; Speigle and Brainard 1996; McCoy et al. 2018, 2019; McCoy and Prum 2019). In the intriguing case of *R. carbo*, velvety red plumage shares its microstructure (and reflectance curve) with super black plumages (Supplemental Data 1); this velvet red, absorptive plumage is adjacent to an extremely bright white bill (Figure 1D). This dark, antireflective frame would thus enhance the perceived brilliance of the “silver” beak, which may play an important role in female choice.

Further evidence for the importance of microstructure in color appearance comes from within-bird plumage variation. Within single birds that have both bright red and dark velvety red plumage (e.g., upper throat versus lower throat in both *R. dimidiatus* and *R. melanogaster*), the relative amount of pigment molecules in bright and dark red patches are highly correlated (Figure 6A,D), suggesting that pigment alone is not responsible for these differences in appearance. By contrast, the feather microstructure is strikingly different between these types of patches (Figure 6B-C,E-F), suggesting that microstructure, not pigment, is responsible for the differences in appearance of bright and dark velvety plumage. These feathers may also have internal nanostructural changes, which have been shown to make carotenoid-based colors brighter (Shawkey and Hill 2005). Research on microstructural variation in colorful displays, including sex differences, is expanding rapidly (Lee et al. 2009; Iskandar et al. 2016; Enbody et al. 2017; McCoy et al. 2018; McCoy and Prum 2019). To gain more insight into evolutionary dynamics, we require a complete understanding of the physical basis of color. This means accounting for the optical effects of microstructure in addition to the traditionally studied pigments and nanostructure (Burns et al. 2017).

### The proxy treadmill: honest signals can be gamed, causing trait elaboration

Why did male feathers evolve elaborate microstructures? If percepts of color influence female choices of mates, whether the percept is an arbitrary preference (Prum 2012) or an index of male quality, then males will be selected to produce this percept in females whether this is by changes to the chemical composition or microstructure of their feathers. From this perspective, the evolutionary dynamics of female preferences and male traits has aspects of an arms race between the conflicting interests of the sexes (Dawkins and Krebs 1978; Krebs and Dawkins 1984; Burk 1988; Hill 1994; Funk and Tallamy 2000; McCoy and Haig 2020). Females establish tests of quality and males are selected to pass these tests whether by ‘honest’ displays or ‘gaming’ the test. Rather than eating and metabolizing more carotenoids to appear more intensely red, males could use microstructural amplifiers to make their red plumage appear brighter and more saturated. Mutant melanic scarlet tanagers (*Piranga olivacea*, Cardinalidae) retain the feather structure typical of carotenoid-containing feathers, suggesting that at least some birds have independent genetic control systems for pigment deposition and feather structure (Brush 1970). Thus, it is possible for selection to act on feather structure without impacting pigment. We suggest that male *Ramphocelus* tanagers may have been selected for structural modifications which enhance their visual appearance and distort the link between appearance and carotenoid content.

Perhaps red carotenoid-based colors were favored by female selection because they were correlated with male quality (Olson and Owens 1998), but selection on males to appear attractive by whatever means gradually diluted the information about male quality conveyed by the male signal (Hill 1994). As males exaggerate their appearance, females are selected to develop additional quality tests to separate the wheat from the chaff (e.g., females may prefer more saturated feathers or additional ornaments). Because females cannot unilaterally abandon a prior quality test without dooming their sons to being unattractive, male ornaments may pile atop one another, potentially causing extreme and elaborate signals. This is the proxy treadmill (McCoy and Haig 2020): signal traits become exaggerated as proxies of quality are continuously modified or replaced (because examinees are under strong selective pressure to inflate their apparent quality, thus devaluing any given proxy).

We are not the first to propose that males may find ways to amplify their carotenoid signals. Male guppies (*Poecilia reticulata*) have orange carotenoid-colored spots for sexual display. Recall that carotenoids must be eaten and metabolized rather than synthesized *de novo*, a commonly-cited reason supporting the idea that carotenoids are honest signals. Guppies synthesize (*de novo*) red pteridine pigments (drosopterins) similar in hue to carotenoids and include these pigments in their red spots. The authors suggest that male guppies use drosopterin pigments “in a manner that dilutes, but does not eliminate, the indicator value of carotenoid coloration” (Grether et al. 2001). Beyond carotenoids, direct evidence for deceptive elements to honest signaling has been reported in many invertebrates (e.g., Wilson et al. 2007; Christy and Rittschof 2010; Ghislandi et al. 2017) and a few vertebrates (Candolin 1999; Bro-Jørgensen and Pangle 2010; Summers 2014); see discussion in (McCoy and Haig 2020).

Further, mate choice is not the only examination that may be susceptible to performance-enhancing strategies. Maternal assessment of embryo quality is another evolutionary arena for signaling and evaluation (McCoy and Haig 2020); embryos are “auditioning” to be chosen and nurtured by the mother (Forsdyke 2019). One measure of embryo quality, the signalling hormone chorionic gonadotropin (CG), has seen an extreme inflation over evolutionary time with no clear functional significance. Human embryos carry 6 copies of the CG gene, have evolved modifications to the molecule that extend CG’s half-life (and thus persistence time as a signal of quality) from ∼21–25 minutes to ∼11-462 hours, and produce “heroic quantities” (Zeleznik 1998) of CG during pregnancy (McCoy and Haig 2020). This extreme elaboration is reminiscent of the microstructural diversity observed in male *Ramphocelus* tanagers. Goodhart’s Law in economics and the analogous Campbell’s Law describe this well-observed phenomenon; pithily phrased, “when a measure becomes a target, it ceases to be a good measure” (Strathern 1997). The classic case is high-stakes testing. When standardized test scores are used as a proxy for how well middle school teachers educate their pupils, determining funding, teachers ‘teach to the test’ rather than improving students’ reading skills (Goldstein 2004). This does not mean tests are useless: for example, the SAT does convey some true information—but this signal is distorted by socioeconomic background, teacher strategy, culture, and test prep programs.

Many alternative explanations could explain our results herein. For example, perhaps microstructures themselves are honest signals. Some research has shown convincingly that nanostructural color is a correlate of individual quality (Keyser and Hill 2000; Hill et al. 2005; Shawkey et al. 2007; White 2020); might the same be true for microstructures? Perhaps beautiful color signals are merely the resolution of arbitrary aesthetic preferences in females (Prum 2012), in which case microstructures that make colors more beautiful are of course preferred. Signals cannot be pigeonholed as purely honest nor purely deceptive, because we are always viewing a snapshot in evolutionary time wherein species are subject to many competing selective pressures. Further work that compares carotenoid presence and microstructural modifications across many males, ideally also with some measure of genetic quality, would be invaluable.

## Author Contributions

All authors conceived the research design. DEM and EH performed all pigment extractions, SEM work, and spectrophotometry. CV and ST designed and implemented the LC-MS procedure for carotenoid characterization. DEM performed the optical simulations. AJS constructed the phylogeny and performed all regressions. DH provided conceptual groundwork. All authors jointly wrote the paper.

## Acknowledgements

We are very grateful to Jeremiah Trimble and Kate Eldridge for their help with the ornithological collections. Special thanks to Dan Utter and Professor Colleen Cavanaugh for their methodological assistance and use of their lab equipment. Thank you to Sofia Prado-Irwin and Dan Utter for helpful comments on the manuscript. Thank you to Gabriel Ugueto for the outstanding artwork, and to Nick Athanas – antpitta.com – for the beautiful photos. This work was performed in part at the Harvard University Center for Nanoscale Systems (CNS), a member of the National Nanotechnology Coordinated Infrastructure Network (NNCI), which is supported by the National Science Foundation under NSF ECCS award no. 1541959. DEM’s research is conducted with Government support under and awarded by DoD, Air Force Office of Scientific Research, National Defense Science and Engineering Graduate (NDSEG) Fellowship, 32 CFR 168a, and DEM and AJS were supported by research funds from the Harvard University Department of Organismic and Evolutionary Biology (OEB).

## Conflict of Interest Statement

The authors declare no conflicts of interest.

## Data Accessibility Statement

All code and data will be made available on Dryad.

## Supplemental Data

- Supplemental Information PDF (tables and figures)
- Supplemental Data S1 (CSV file)

## Notes

### Competing Interest Statement

The authors have declared no competing interest.

