## Supplemental Information for "Microstructures amplify carotenoid plumage signals in colorful tanagers"

**Supplemental Materials in this PDF:**

**Additional Supplemental Materials:**

**Supplementary Data 1:** Complete Spectrophotometry Results.

- CSV file entitled "Supplementary_Table_Spectrophotometry_Results.csv" includes complete spectrophotometry results for all 10 species, including 6 body regions per species measured both at 90° and 45° incident light.

**Figure S1: Ancestral state reconstructions of each carotenoid pigment family.**

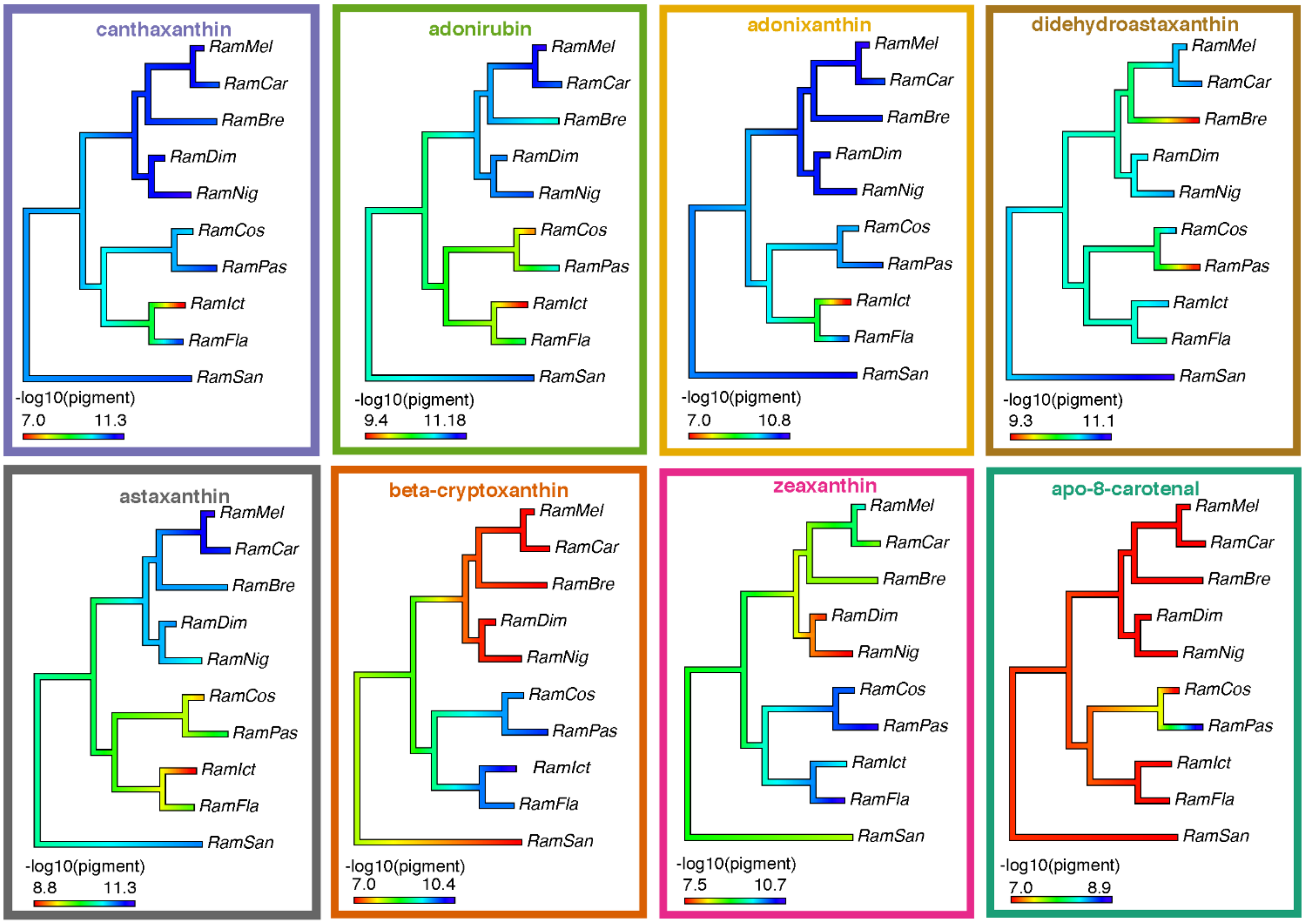

**Figure S1: Ancestral state reconstructions of each carotenoid pigment family.** We summed and log-transformed all isoforms within each pigment family, and then estimated ancestral states for each pigment.

**Figure S2: Metabolic map of carotenoid pigments.**

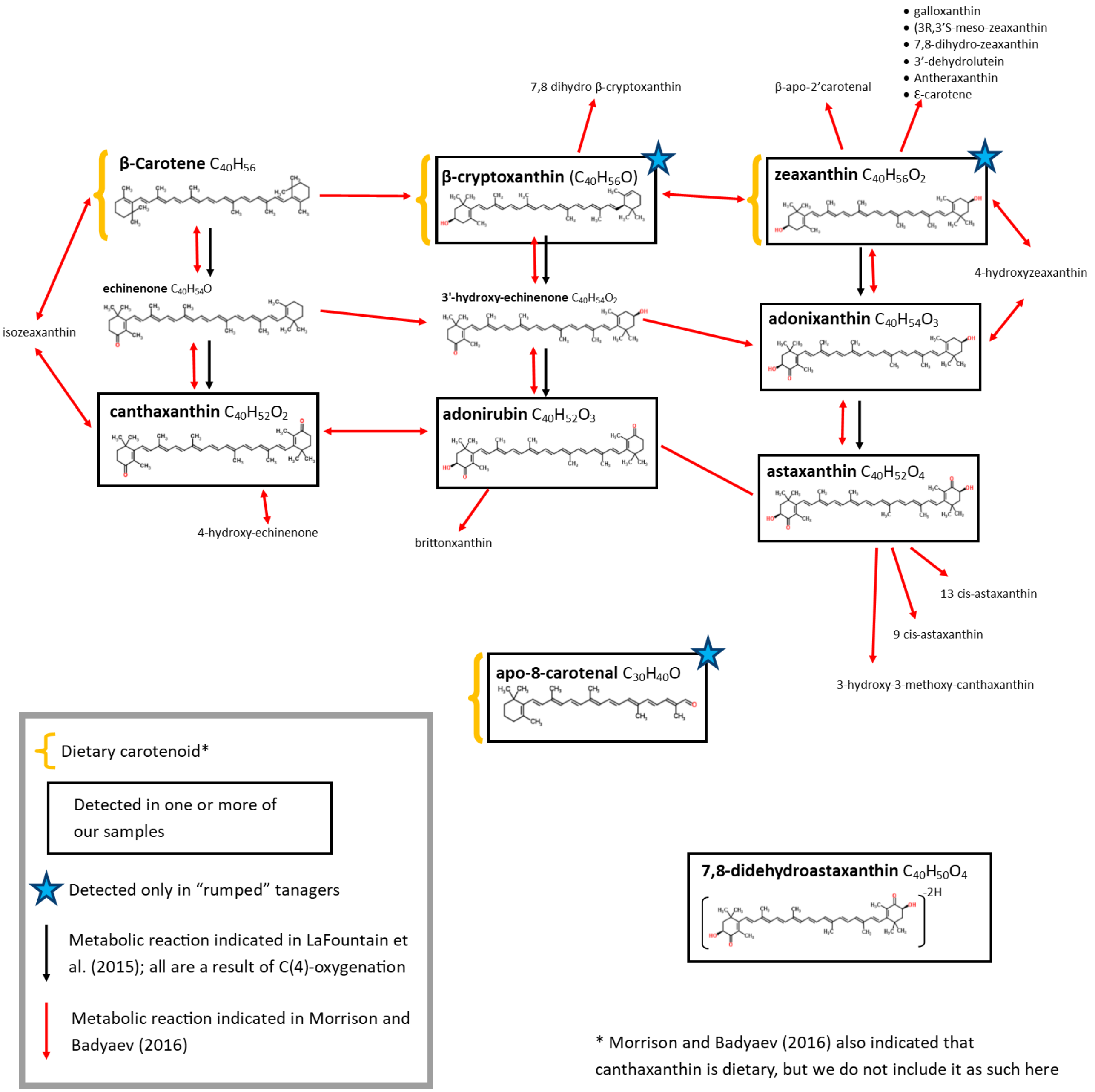

**Figure S2: Metabolic map of carotenoid pigments.** Based upon metabolic maps in the literature, we mapped our detected pigment families (in black boxes) into a metabolic network to demonstrate the presumed relationships between identified pigments. Apo-8-carotenal and didehydroastaxanthin could not be incorporated into the map and thus are shown separately.

**Figure S3: LC-MS pigment characterization is repeatable across individuals within a species.**

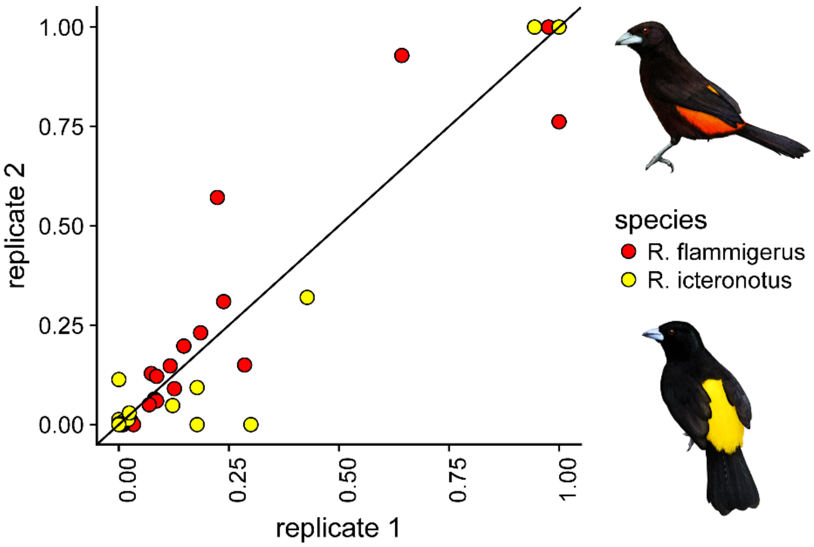

**Figure S3**: **LC-MS pigment characterization is consistent across individuals within a species.** Pigment profiles for 2 individuals per species of *R. f. icteronorus* and *R. flammigerus* demonstrate significant within-species correlations. All values are normalized within an individual by feather weight, and normalized such that the maximum signal was set to 1. Linear regression output for *R. flammigerus*: slope = 0.88, SE = 0.066, $R^2$ = 0.87, $p < 0.0005$. Linear regression output for *R. f. icteronotus*: slope = 0.96, SE = 0.054, $R^2$ = 0.92, $p < 0.0005$. Artwork in bird silhouettes credit Gabriel Ugueto.

**Figure S4: Additional PCAs uphold main results.**

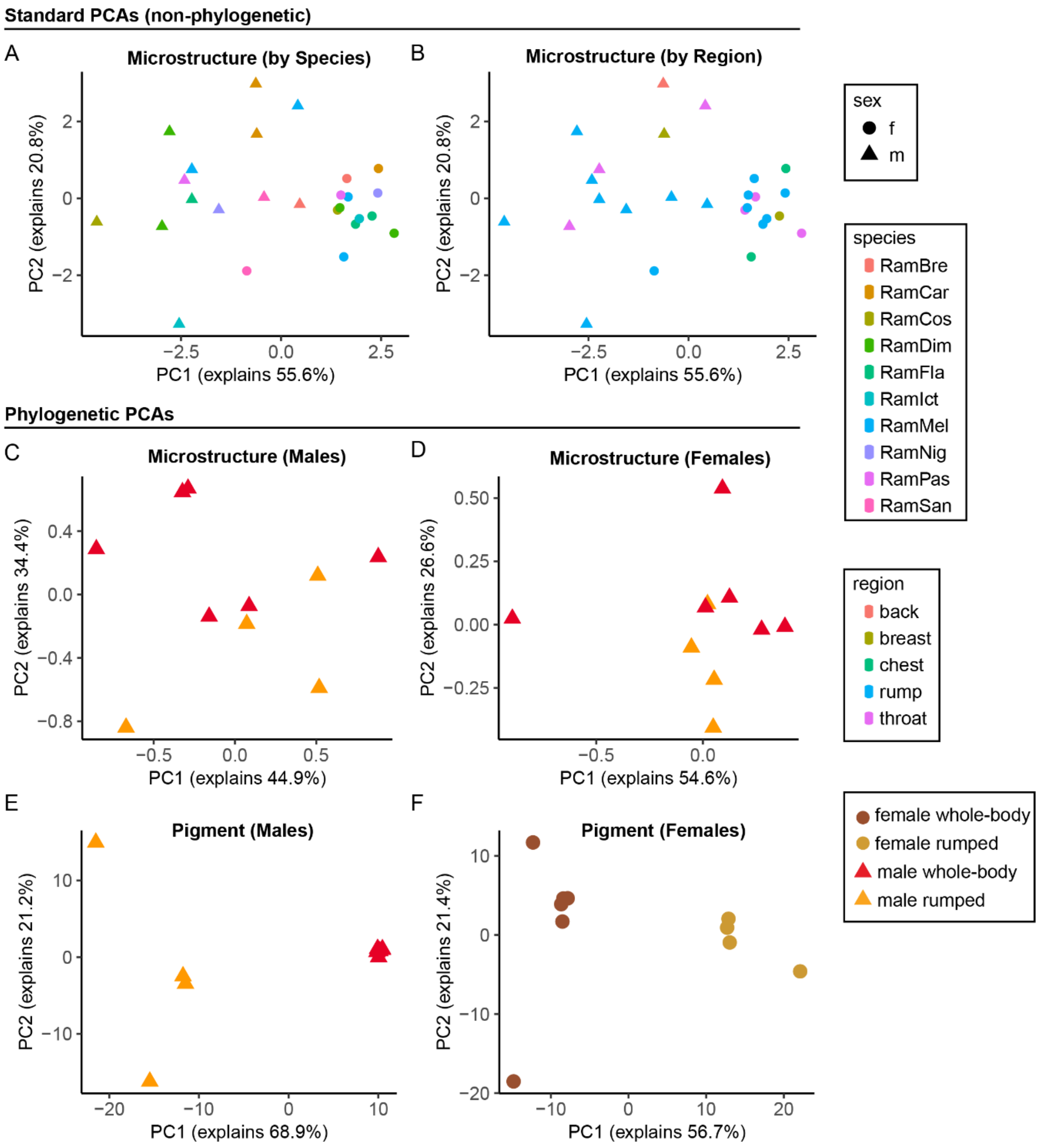

**Figure S4: Additional PCAs uphold main results.** A: Non-phylogenetic PCA of microstructural measurements for all measured feathers, males and females, grouped by species. B: Non-phylogenetic PCA of all microstructural measurements from all measured feathers grouped by region. C. Phylogenetic PCA of microstructural measurements for males (one feather per male per species). D: Phylogenetic PCA of microstructural measurements for females (one feather per female per species). E: Phylogenetic PCA of pigment families for males (one feather per male per species). F: Phylogenetic PCA of pigment families for females (one feather per female per species). For species where we had to select only one patch, we chose the following: Females: *R. melanogas*ter throat, *R. dimidiatus* rump, *R. flammigerus* rump, *R. passerinii* rump. Males: *R. dimidiatus* rump, *R. carbo* back.

**Figure S5: Complete SEM Results**

A. *Ramphocelus bresilius*

Female rump

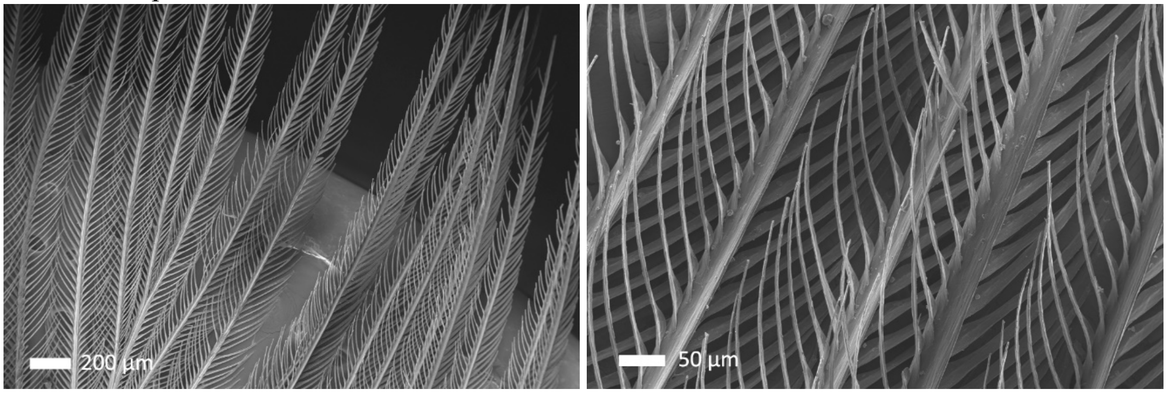

Male rump

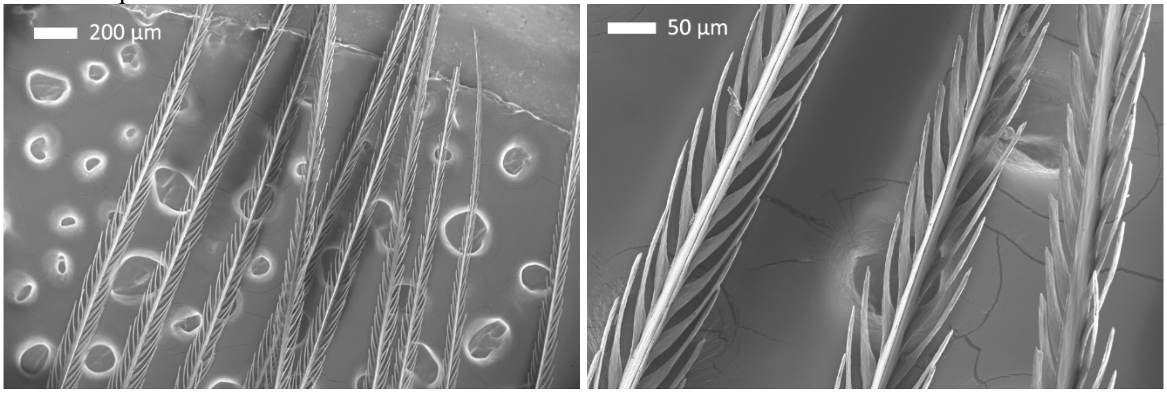

B. *Ramphocelus carbo*

Female chest

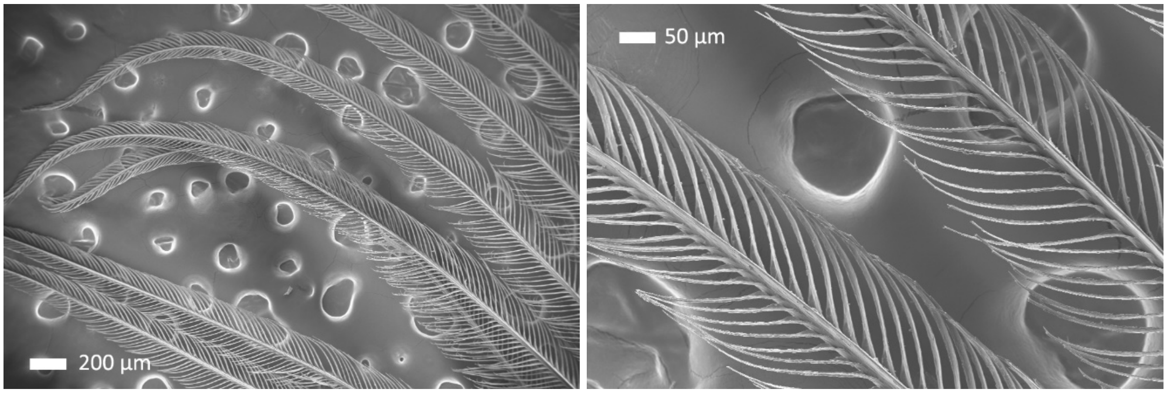

Male velvet red back:

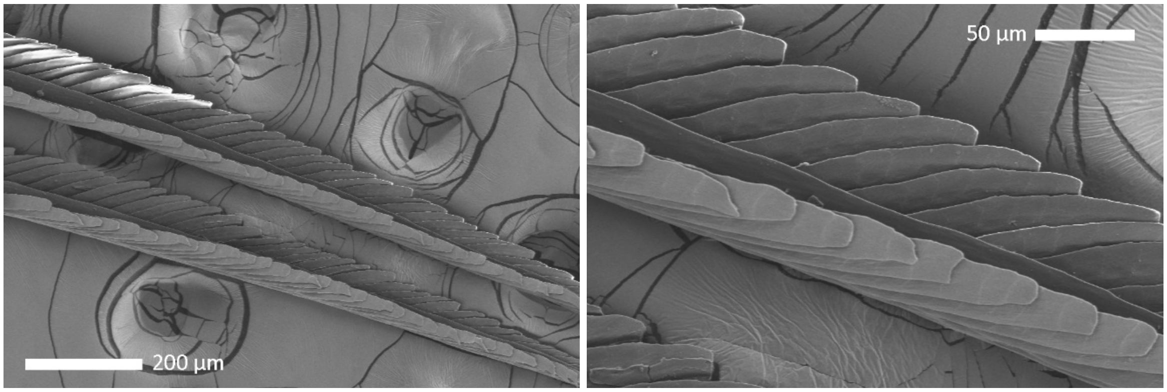

Red breast:

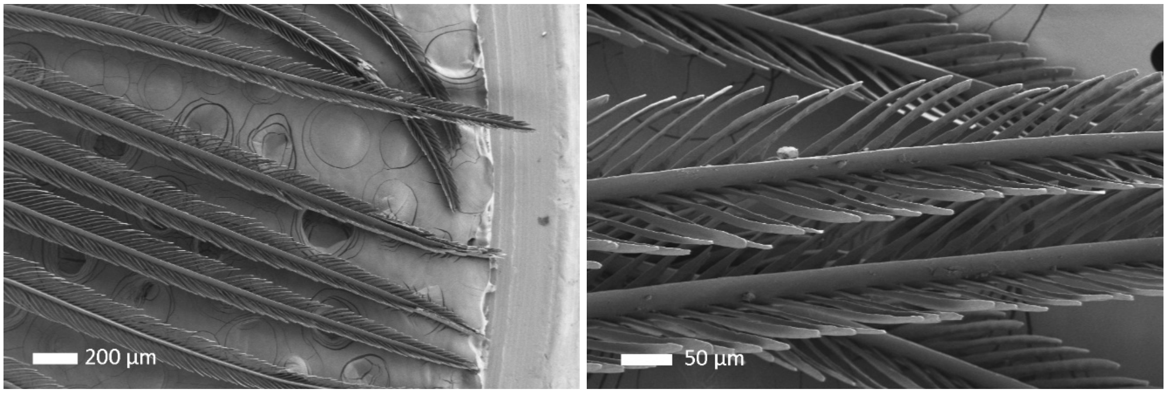

C. *Ramphocelus passerinii costaricensis*

Female throat

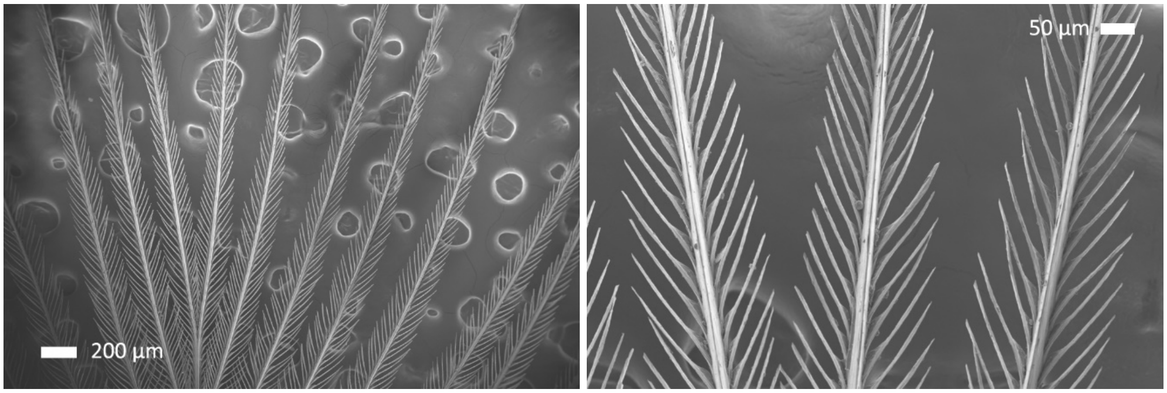

Male rump

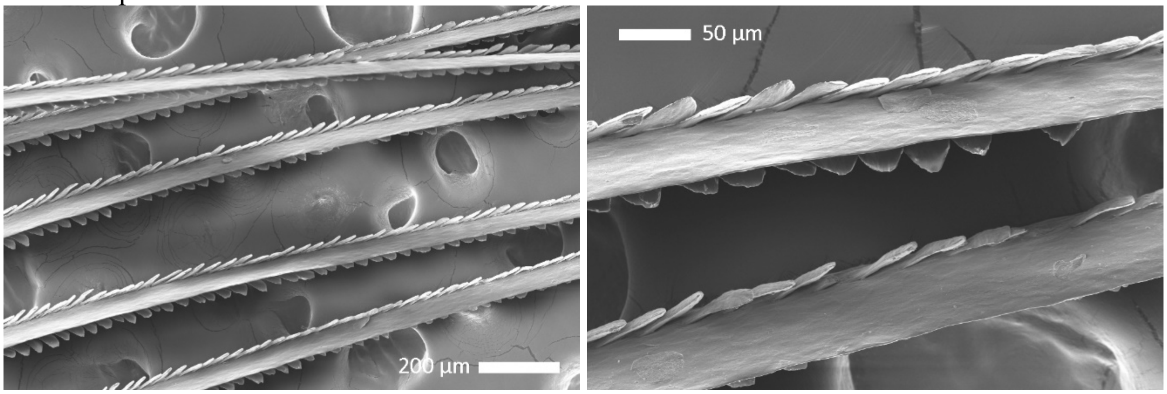

D. *Ramphocelus dimidiatus*

Female rump:

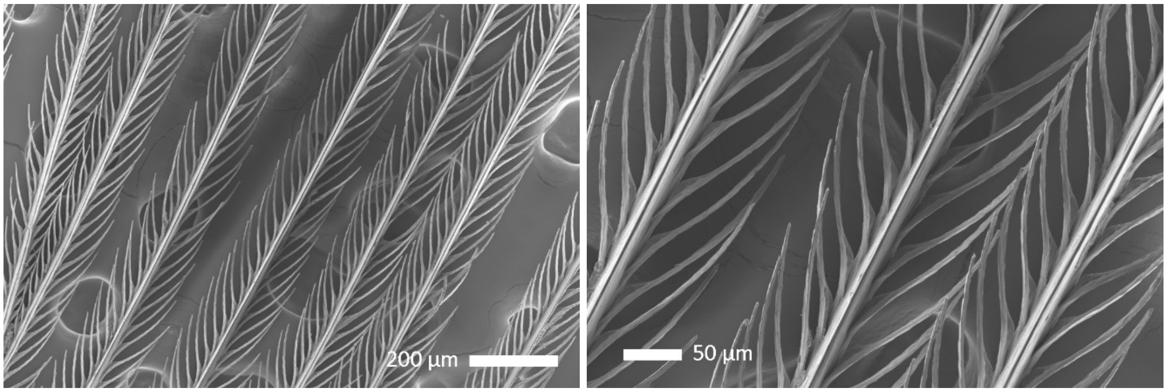

Female Throat:

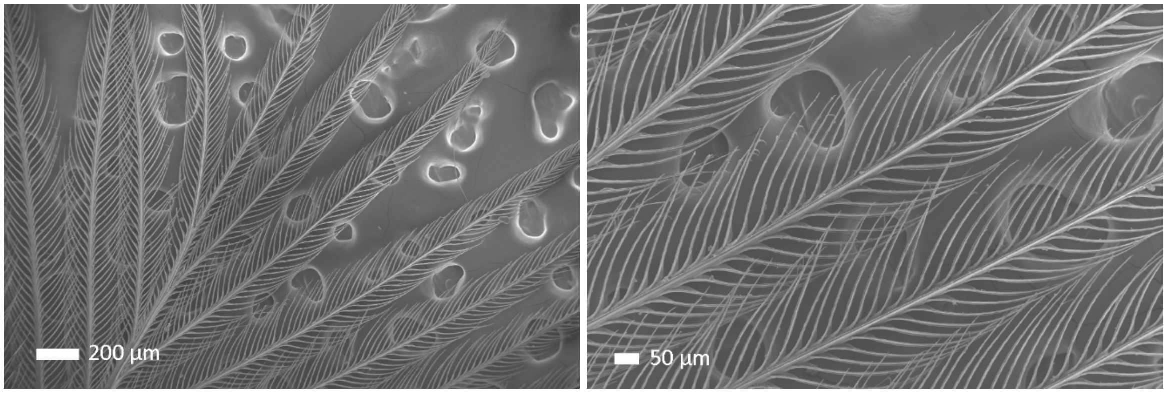

Male rump:

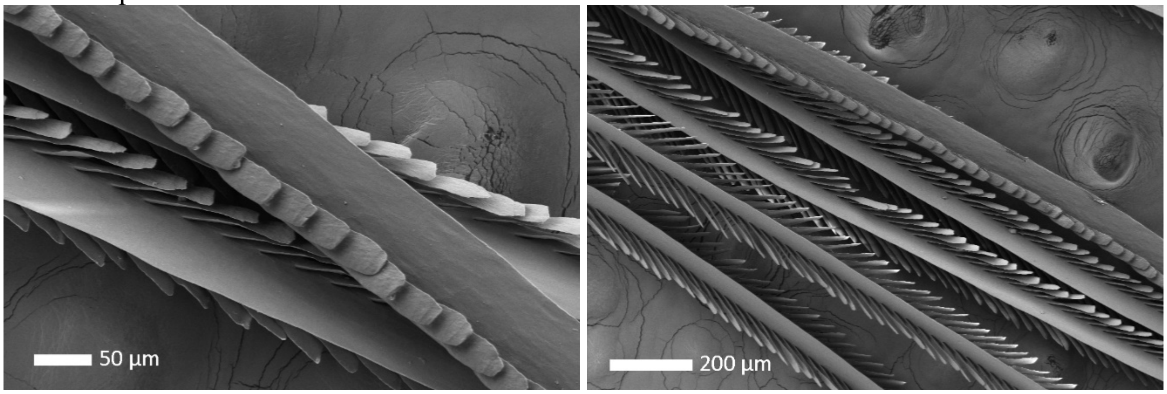

Male throat:

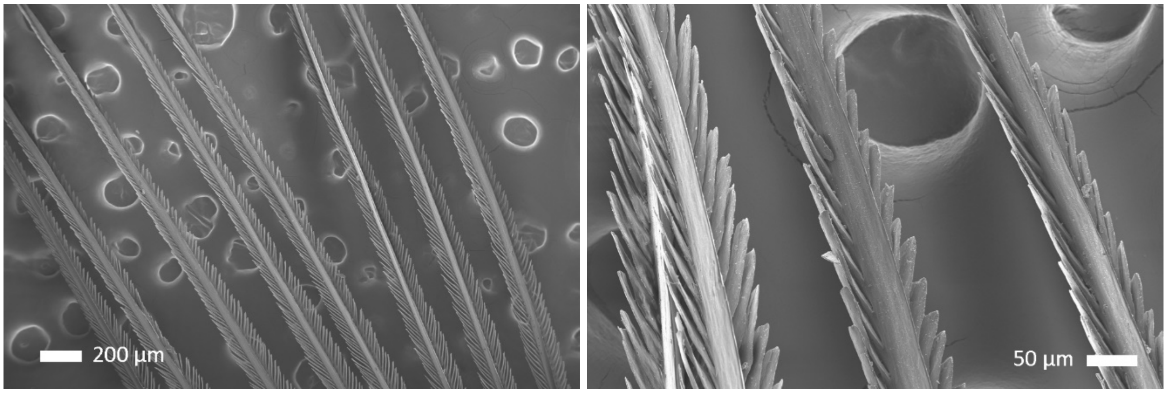

E. *Ramphocelus flammigerus*

Female breast:

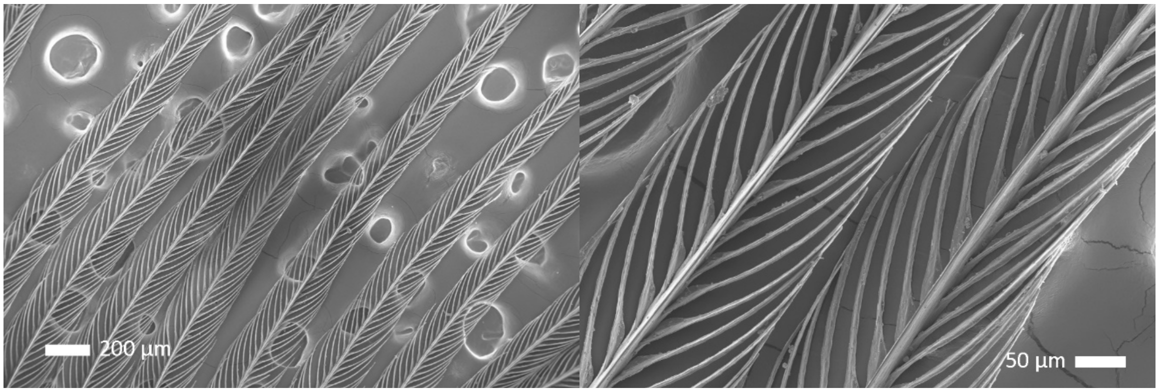

Female rump:

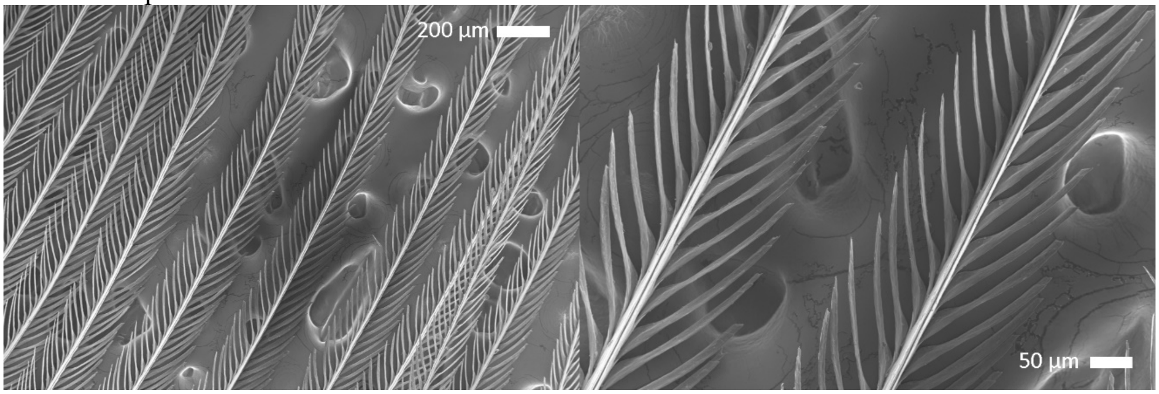

Male rump

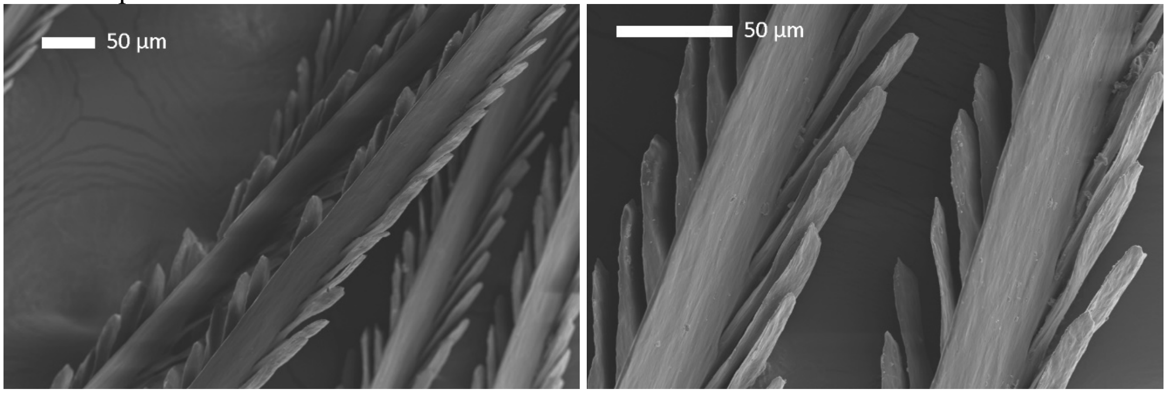

F. *Ramphocelus flammigerus icteronotus*

Female rump

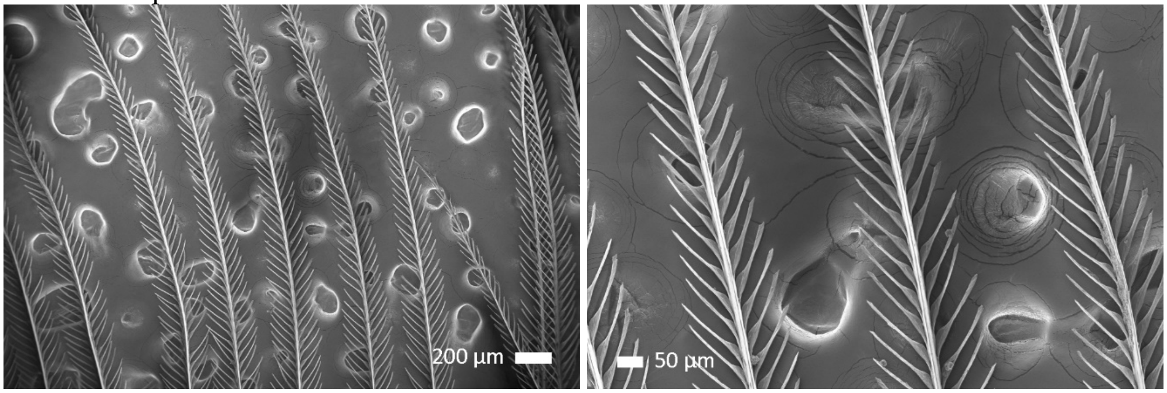

Male rump

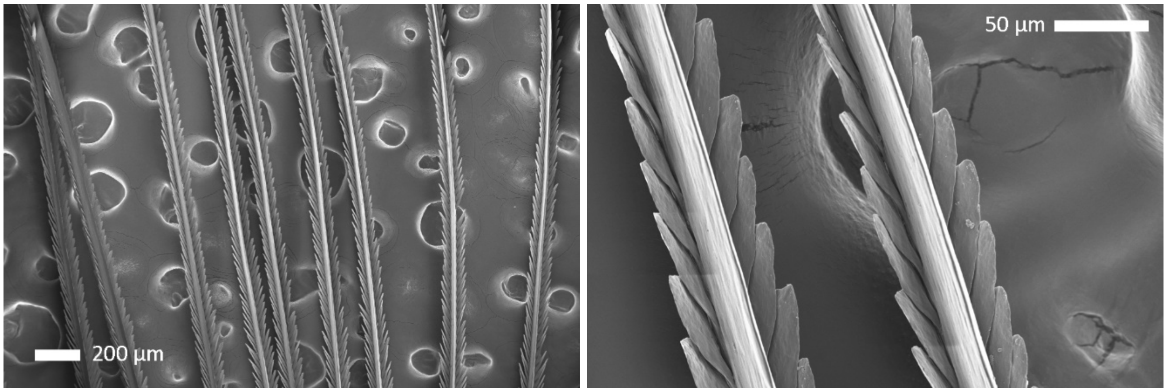

G. *Ramphocelus melanogaster*

Female chest

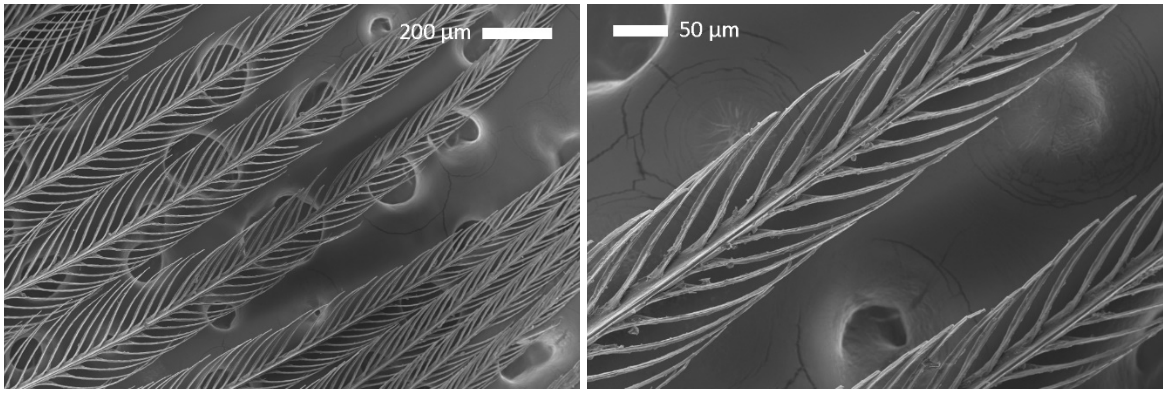

Male Throat Upper:

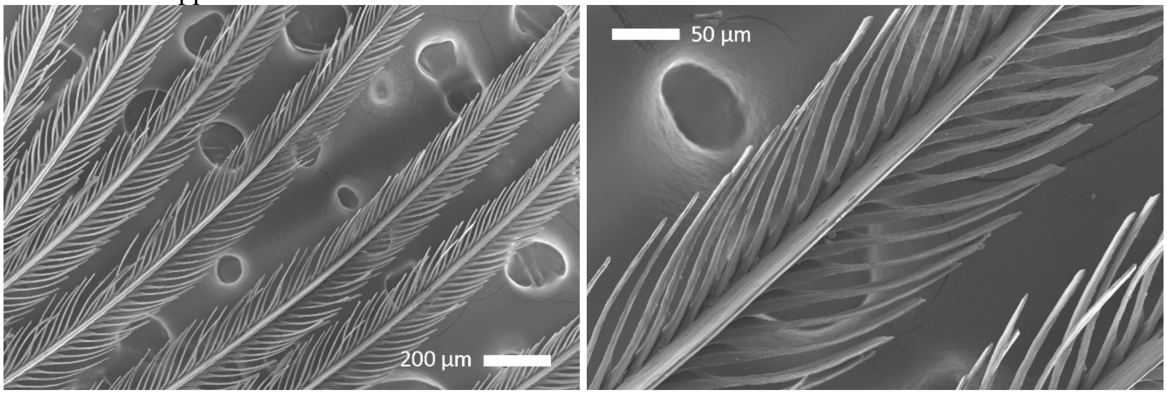

Male Throat Lower:

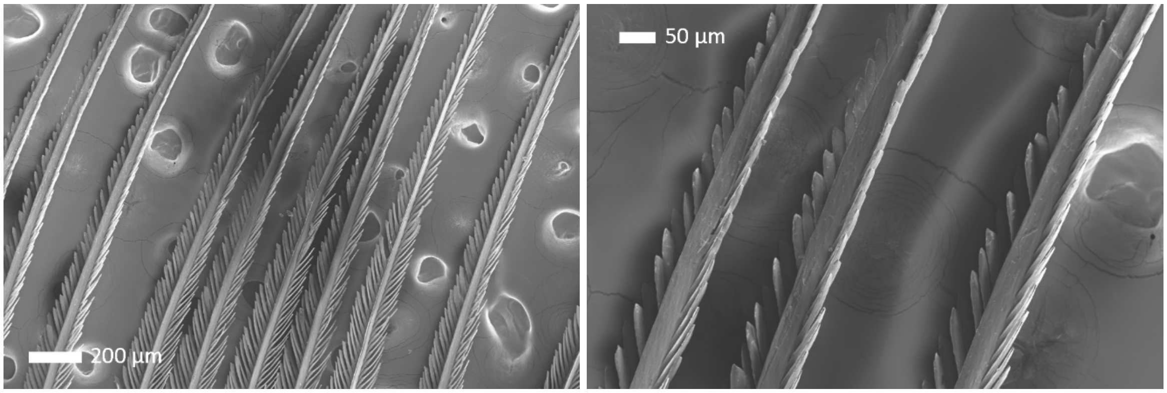

H. *Ramphocelus nigrogularis*

Female rump

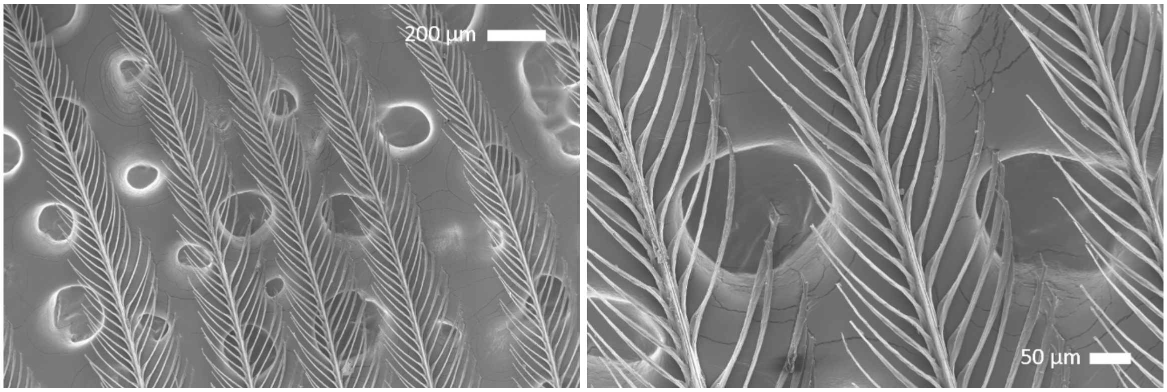

Male rump

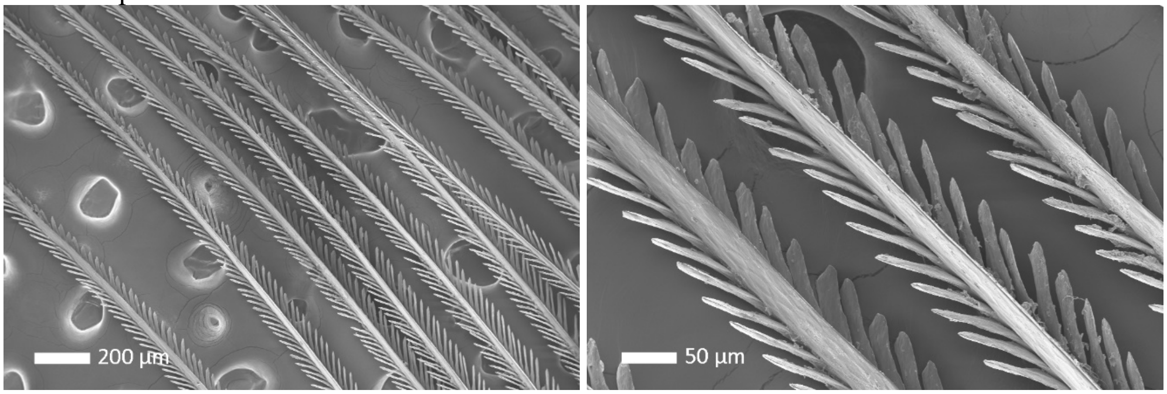

I. *Ramphocelus passerinii*

Female rump

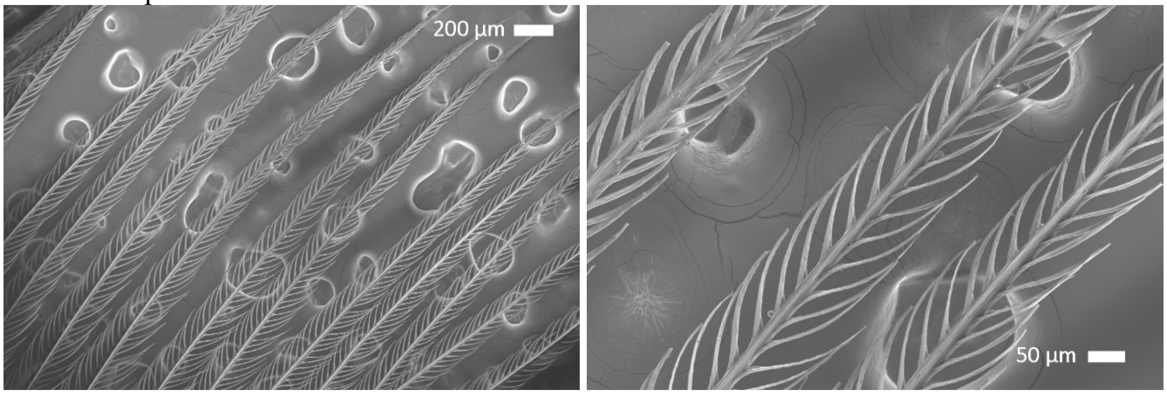

Male rump

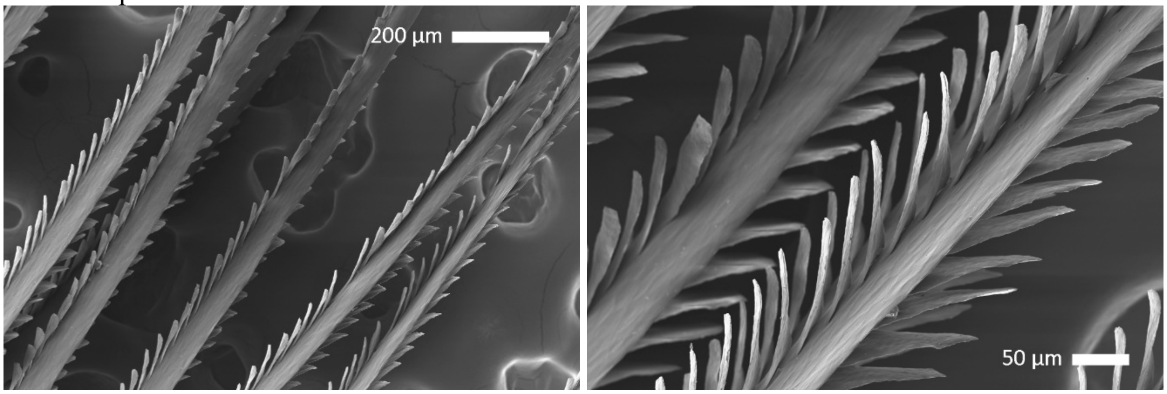

J. *Ramphocelus sanguinolentus*

Female rump

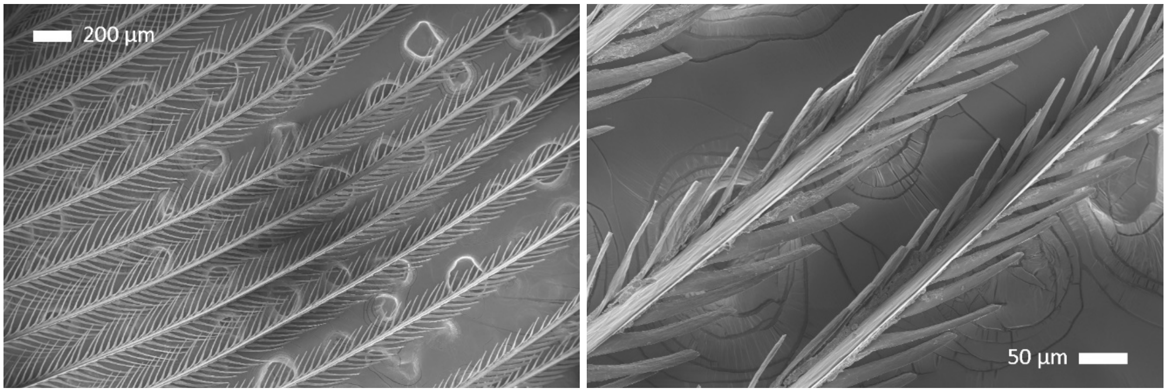

Male rump

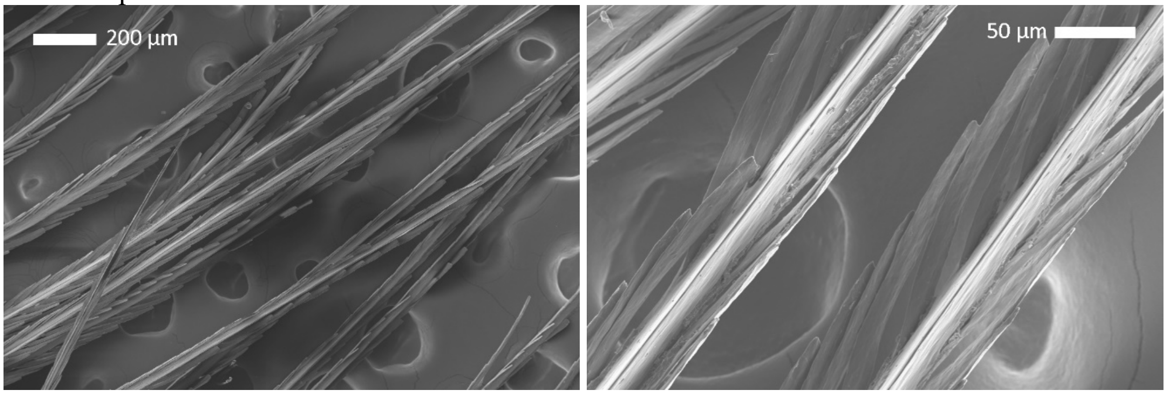

**Table S1: Specimen details.**

| Scientific Name | Sex | Specimen Number | Feather Region | Qualitative Patch Color |
|---|---|---|---|---|
| *R. sanguinolentus* | M | MCZ 110057 | rump | red |
| *R. sanguinolentus* | F | MCZ 11055 | rump | red |
| *R. nigrogularis* | M | MCZ 139568 | rump | red |
| *R. nigrogularis* | F | MCZ 299482 | rump | red |
| *R. dimidiatus** | M | MCZ 105428, 105430, 105431 | throat | dark velvet red |
| *R. dimidiatus** | M | MCZ 105428, 105430, 105431 | rump | bright red |
| *R. dimidiatus* | F | MCZ 105445 | throat | dark red |
| *R. dimidiatus* | F | MCZ 105445 | rump | red |
| *R. melanogaster* | M | MCZ 96403 | throat (upper) | dark velvet red |
| *R. melanogaster* | M | MCZ 96403 | throat (lower) | bright red |
| *R. melanogaster* | F | MCZ 96404 | chest | dark red |
| *R. melanogaster* | F | MCZ 96404 | throat | red |
| *R. carbo** | M | MCZ 299748, 299750, 299751 | breast | dark red |
| *R. carbo* | M | MCZ 299751 | back | dark red-black velvet |
| *R. carbo* | F | MCZ 299749 | chest | dark red |
| *R. bresilius* | M | MCZ 273881 | rump | red |
| *R. bresilius* | F | MCZ 273880 | rump | red |
| *R. costaricensis* | M | MCZ 108121 | rump | red-orange |
| *R. costaricensis* | F | MCZ 108141 | throat | orange-yellow |
| *R. costaricensis* | F | MCZ 108141 | rump | orange |
| *R. passerinii* | M | MCZ 110096 | rump | red |
| *R. passerinii* | F | MCZ 110103 | rump | reddish-orange |
| *R. flammigerus* | M | MCZ 103856 | rump | orange |
| *R. flammigerus* | F | MCZ 103892 | breast | reddish-orange |
| *R. flammigerus* | F | MCZ 103892 | rump | orange-yellow |
| *R. flammigerus icteronotus* | M | MCZ 107435 | rump | yellow |
| *R. flammigerus icteronotus* | F | MCZ 107448 | rump | yellow |

**Table S1: Specimen Details.** Specimens used in pigment extractions and SEM imaging. * indicates species for whom multiple individuals were combined for pigment extraction (to determine minimum amount of feather necessary). MCZ refers to the Harvard Museum of Comparative Zoology.

**Table S2: NCBI Accession Numbers.**

| Species | NCBI Accession Number |
|---|---|
| *Ramphocelus carbo* | U15723 |
| *Ramphocelus melanogaster* | FJ799883 |
| *Ramphocelus bresilius* | U15724 |
| *Ramphocelus dimidiatus* | FJ799881 |
| *Ramphocelus nigrogularis* | U15721 |
| *Ramphocelus costaricensis* | U15722 |
| *Ramphocelus passerinii* | EF529965 |
| *Ramphocelus flammigerus* | KR817429 |
| *Ramphocelus f. icteronotus* | U15719 |
| *Ramphocelus sanguinolentus* | U15718 |
| *Tachyphonus coronatus* | FJ799885 |
| *Tachyphonus rufus* | FJ799896 |
| *Tachyphonus phoenicius* | FJ799893 |
| *Eucometis penicillata* | FJ799875 |
| *Lanio fulvus* | EU647917 |
| *Tachyphonus luctuosus* | EF529967 |

**Table S3: Microstructural measurements**

| species | sex | region | barb width (maximum) | | barb width (top-down) | | barbule width | | inter-barbule distance (for 5 barbules) | | barb-barbule angle | | barbule length | |
|---|---|---|---|---|---|---|---|---|---|---|---|---|---|---|
| | | | mean | sd | mean | sd | mean | sd | mean | sd | mean | sd | mean | sd |
| *R. bresilius* | f | rump | 30.78 | 2.71 | 26.30 | 1.42 | 5.35 | 0.78 | 195.89 | 11.16 | 31.10 | 3.36 | 227.29 | 15.39 |
| *R. carbo* | f | chest | 21.01 | 1.83 | 16.33 | 2.14 | 6.63 | 1.36 | 175.67 | 12.02 | 39.95 | 3.85 | 232.95 | 23.18 |
| *R. costaricensis* | f | throat | 21.96 | 2.69 | 20.45 | 1.16 | 5.88 | 1.41 | 200.95 | 12.98 | 27.52 | 2.94 | 167.01 | 6.55 |
| *R. dimidatus* | f | rump | 22.76 | 2.30 | 20.61 | 1.36 | 5.83 | 1.10 | 213.06 | 23.63 | 28.99 | 4.71 | 161.36 | 13.38 |
| *R. dimidatus* | f | throat | 20.00 | 2.97 | 16.55 | 2.76 | 6.50 | 1.98 | 227.87 | 15.98 | 35.87 | 3.64 | 284.49 | 25.91 |
| *R. flammigerus* | f | breast | 18.17 | 1.45 | 16.74 | 2.29 | 6.77 | 2.13 | 230.08 | 26.58 | 29.05 | 3.99 | 227.35 | 11.58 |
| *R. flammigerus* | f | rump | 18.22 | 1.14 | 18.14 | 2.89 | 7.70 | 2.22 | 236.94 | 12.77 | 31.37 | 4.86 | 191.65 | 6.88 |
| *R. f. icteronotus* | f | rump | 18.22 | 0.86 | 16.19 | 1.90 | 7.27 | 2.33 | 263.52 | 7.25 | 35.79 | 5.92 | 146.51 | 18.61 |
| *R. melanogaster* | f | chest | 19.76 | 1.30 | 17.08 | 2.07 | 5.94 | 1.43 | 243.24 | 9.65 | 22.43 | 1.09 | 171.70 | 12.09 |
| *R. melanogaster* | f | throat | 22.31 | 2.49 | 20.67 | 2.08 | 6.50 | 1.14 | 189.69 | 5.93 | 32.66 | 1.73 | 196.60 | 27.31 |
| *R. nigrogularis* | f | rump | 19.89 | 1.62 | 15.51 | 0.96 | 5.78 | 2.24 | 208.80 | 16.61 | 28.14 | 2.86 | 232.46 | 25.80 |
| *R. passerinii* | f | rump | 23.17 | 4.17 | 17.96 | 1.28 | 6.80 | 1.47 | 230.51 | 6.59 | 30.15 | 2.52 | 157.49 | 22.09 |
| *R. sanguinolentis* | f | rump | 31.70 | 4.47 | 23.22 | 3.72 | 12.14 | 2.39 | 249.25 | 20.78 | 18.40 | 3.91 | 136.49 | 22.73 |
| *R. bresilius* | m | rump | 22.29 | 1.83 | 15.25 | 1.21 | 9.97 | 1.31 | 258.94 | 17.94 | 22.66 | 5.87 | 121.23 | 13.79 |
| *R. carbo* | m | back | 42.89 | 4.15 | 14.85 | 1.77 | 17.45 | 0.79 | 169.31 | 10.59 | 24.97 | 11.17 | 162.93 | 11.65 |
| *R. carbo* | m | breast | 36.59 | 1.91 | 19.76 | 2.50 | 13.40 | 2.41 | 179.82 | 11.91 | 29.08 | 9.00 | 124.68 | 17.29 |
| *R. costaricensis* | m | rump | 43.45 | 1.04 | 42.26 | 3.40 | 24.80 | 3.15 | 176.38 | 12.49 | 15.16 | 4.29 | 60.31 | 4.94 |
| *R. dimidatus* | m | throat | 32.98 | 3.00 | 32.37 | 10.63 | 13.79 | 3.23 | 163.03 | 11.11 | 14.38 | 1.66 | 78.85 | 14.92 |
| *R. dimidiatus* | m | rump | 81.43 | 6.27 | 45.36 | 5.30 | 13.41 | 1.85 | 213.76 | 5.14 | 20.48 | 3.94 | 101.00 | 13.24 |
| *R. flammigerus* | m | rump | 36.00 | 1.62 | 32.50 | 0.85 | 11.91 | 2.02 | 164.69 | 6.36 | 19.28 | 6.66 | 87.66 | 9.67 |
| *R. f. icteronotus* | m | rump | 27.85 | 2.05 | 14.00 | 9.81 | 16.24 | 3.21 | 174.24 | 8.09 | 15.88 | 5.42 | 63.79 | 2.98 |
| *R. melanogaster* | m | throat | 43.69 | 3.11 | 21.25 | 1.55 | 7.82 | 1.77 | 158.56 | 11.94 | 30.03 | 6.96 | 175.78 | 8.49 |
| *R. melanogaster* | m | throat | 39.35 | 0.98 | 27.92 | 1.35 | 15.35 | 1.29 | 172.80 | 19.30 | 22.81 | 2.74 | 85.85 | 16.76 |
| *R. nigrogularis* | m | rump | 29.45 | 1.06 | 28.45 | 2.17 | 12.93 | 3.80 | 185.56 | 11.12 | 25.08 | 2.17 | 87.65 | 5.79 |
| *R. passerinii* | m | rump | 49.22 | 6.93 | 43.67 | 6.87 | 19.06 | 1.57 | 205.95 | 8.88 | 26.82 | 4.23 | 110.98 | 9.52 |
| *R. sanguinolentis* | m | rump | 32.84 | 1.72 | 22.18 | 2.46 | 18.98 | 0.61 | 327.04 | 19.86 | 19.07 | 3.51 | 196.91 | 11.82 |

**Table S3: Microstructural measurements.** Measurements of SEM photos for all feathers from all species in µm.

**Table S4: Pigment identification using LC-MS.**

| pigment family | molecule name | Monoisotopic mass | formula | retention time (min) | identification |
|---|---|---|---|---|---|
| apo-8-carotenal | apo-8-carotenal | 416.32 | $C_{30}H_{40}O$ | 4.8 | matched pigment standard |
| β-Cryptoxanthin | β -Cryptoxanthin_1 | 552.43 | $C_{40}H_{56}O$ | 3.8 | inferred |
| | β -Cryptoxanthin_2 | 552.43 | $C_{40}H_{56}O$ | 4.1 | inferred |
| | β -Cryptoxanthin_3 | 552.43 | $C_{40}H_{56}O$ | 4.4 | inferred |
| | β -Cryptoxanthin_4 | 552.43 | $C_{40}H_{56}O$ | 4.8 | inferred |
| | β -Cryptoxanthin_5 | 552.43 | $C_{40}H_{56}O$ | 5.7 | inferred |
| canthaxanthin | canthaxanthin_1 | 564.40 | $C_{40}H_{52}O_2$ | 3.15 | possible isomer |
| | canthaxanthin_2 | 564.40 | $C_{40}H_{52}O_2$ | 3.4 | possible isomer |
| | canthaxanthin_3 | 564.40 | $C_{40}H_{52}O_2$ | 3.7 | possible isomer |
| | canthaxanthin | 564.40 | $C_{40}H_{52}O_2$ | 4.3 | matched pigment standard |
| | canthaxanthin.isomer | 564.40 | $C_{40}H_{52}O_2$ | 4.8 | isomer of pigment standard |
| zeaxanthin | zeaxanthin_1 | 568.43 | $C_{40}H_{56}O_2$ | 3.6 | inferred |
| | zeaxanthin_2 | 568.43 | $C_{40}H_{56}O_2$ | 3.8 | inferred |
| | zeaxanthin_3 | 568.43 | $C_{40}H_{56}O_2$ | 4.6 | inferred |
| | zeaxanthin_4 | 568.43 | $C_{40}H_{56}O_2$ | 5 | inferred |
| | zeaxanthin_5 | 568.43 | $C_{40}H_{56}O_2$ | 5.3 | inferred |
| | zeaxanthin_6 | 568.43 | $C_{40}H_{56}O_2$ | 6.3 | inferred |
| adonirubin | adonirubin_1 | 580.40 | $C_{40}H_{52}O_3$ | 2.9 | inferred |
| | adonirubin_2 | 580.40 | $C_{40}H_{52}O_3$ | 3.4 | inferred |
| | adonirubin_3 | 580.40 | $C_{40}H_{52}O_3$ | 3.8 | inferred |
| | adonirubin_4 | 580.40 | $C_{40}H_{52}O_3$ | 4.6 | inferred |
| adonixanthin | adonixanthin_1 | 582.41 | $C_{40}H_{54}O_3$ | 3.2 | inferred |
| | adonixanthin_2 | 582.41 | $C_{40}H_{54}O_3$ | 3.4 | inferred |
| | adonixanthin_3 | 582.41 | $C_{40}H_{54}O_3$ | 3.7 | inferred |
| | adonixanthin_4 | 582.41 | $C_{40}H_{54}O_3$ | 4.31 | inferred |
| didehydroastaxanthin | didehydroastaxanthin | | $C_{40}H_{50}O_4$ | 2.5 | presumed |
| Astaxanthin | astaxanthin | 596.39 | $C_{40}H_{52}O_4$ | 2.8 | matched pigment standard |
| | astaxanthin_1 | 596.39 | $C_{40}H_{52}O_4$ | 3.1 | possible isomer |
| | astaxanthin.isomer | 596.39 | $C_{40}H_{52}O_4$ | 3.3 | isomer of pigment standard |

**Table S4: Pigment identification using LC-MS.** All of the carotenoid molecules found in *Ramphocelus* tanagers were identified in one of three ways (see "identification" column). "Matched pigment standard" indicates that the molecule matched a pigment standard run simultaneously with the samples. "Isomer of pigment standard" indicates that the molecule was identifiable as an isomer of that pigment standard. "Possible isomer" indicates that it is likely to be an isomer of the pigment standard based on similar retention time and MS/MS spectra. "Inferred" indicates a standard was not available for the family, and the identity was inferred based on accurate mass, retention time, MS/MS spectra comparison with libraries, and pigments commonly found in bird feathers as described in the literature.

**Table S5: PCA Loadings for microstructure PCAs (normal and phylogenetic).**

| | Normal PCA (all) | | PhyloPCA (males) | | PhyloPCA (females) | |
|---|---|---|---|---|---|---|
| | PC1 | PC2 | PC1 | PC2 | PC1 | PC2 |
| barb width (maximum) | -0.31 | 0.53 | 0.881427 | 0.44595 | -0.63566 | 0.727779 |
| barb width (top-down) | -0.39 | -0.10 | 0.837797 | -0.45126 | -0.50981 | 0.7588 |
| barbule width | -0.42 | 0.06 | 0.299406 | -0.14884 | -0.83465 | -0.36525 |
| inter-barbule distance | 0.19 | -0.24 | -0.19279 | 0.591586 | -0.57043 | -0.63815 |
| barb-barbule angle | 0.39 | 0.26 | 0.013186 | 0.602456 | 0.873479 | -0.19204 |
| barbule length | 0.43 | 0.15 | -0.2042 | 0.950605 | 0.725528 | 0.418 |
| Barb oblongness (top-down/maximum) | 0.01 | 0.74 | | | | |
| Barbule width over length | -0.46 | -0.05 | | | | |

**Table S6: PCA Loadings for pigment PCAs (normal and phylogenetic).**

Normal PCA

| | PC1 | PC2 |
|---|---|---|
| apo-8-carotenal | -0.036 | 0.19 |
| beta-cryptoxanthin*_1 | -0.30 | 0.15 |
| beta-cryptoxanthin*_2 | -0.27 | 0.20 |
| beta-cryptoxanthin*_3 | -0.26 | 0.07 |
| beta-cryptoxanthin*_4 | -0.22 | 0.24 |
| beta-cryptoxanthin*_5 | -0.23 | 0.19 |
| canthaxanthin*_1 | 0.22 | 0.24 |
| canthaxanthin*_2 | 0.17 | 0.27 |
| canthaxanthin*_3 | 0.21 | 0.20 |
| canthaxanthin | 0.21 | 0.21 |
| canthaxanthin_iso | 0.20 | 0.25 |
| zeaxanthin*_1 | -0.18 | 0.24 |
| zeaxanthin*_2 | -0.22 | 0.14 |
| zeaxanthin*_3 | -0.24 | 0.15 |
| zeaxanthin*_4 | -0.11 | 0.30 |
| zeaxanthin*_5 | -0.032 | 0.16 |
| zeaxanthin*_6 | 0.0034 | 0.14 |
| adonirubin*_1 | 0.17 | 0.22 |
| adonirubin*_2 | 0.24 | 0.08 |
| adonirubin*_3 | 0.20 | -0.035 |
| adonirubin*_4 | 0.040 | -0.14 |
| adonixanthin*_1 | 0.21 | 0.24 |
| adonixanthin*_2 | 0.082 | 0.024 |
| adonixanthin*_3 | 0.11 | 0.31 |
| adonixanthin*_4 | 0.0084 | -0.0025 |
| didehydroastaxanthin* | -0.03 | 0.052 |
| astaxanthin | 0.25 | 0.16 |
| astaxanthin*_1 | 0.18 | -0.13 |
| astaxanthin_iso | 0.15 | -0.11 |

Phylogenetic PCA

| | males | | females | |
|---|---|---|---|---|
| | PC1 | PC2 | PC1 | PC2 |
| apo-8-carotenal | -0.38 | -0.72 | | |
| cryptoxanthin | -0.98 | -0.12 | 0.97 | -0.07 |
| cantaxanthin | 0.64 | -0.35 | -0.71 | 0.018 |
| zeaxanthin | -0.80 | -0.30 | 0.50 | -0.38 |
| adonirubin | 0.85 | -0.36 | -0.67 | 0.18 |
| adonixanthin | 0.63 | -0.72 | -0.47 | -0.28 |
| didehydroastaxanthin | 0.22 | 0.50 | 0.39 | 0.91 |
| astaxanthin | 0.92 | -0.26 | -0.96 | 0.09 |
